# The crystal structure of SnTox3 from the necrotrophic fungus *Parastagonospora nodorum* reveals a unique effector fold and insights into Kex2 protease processing of fungal effectors

**DOI:** 10.1101/2020.05.27.120113

**Authors:** Megan A. Outram, Yi-Chang Sung, Daniel Yu, Bayantes Dagvadorj, Sharmin A. Rima, David A. Jones, Daniel J. Ericsson, Jana Sperschneider, Peter S. Solomon, Bostjan Kobe, Simon J. Williams

## Abstract

- Plant pathogens cause disease through secreted effector proteins, which act to modulate host physiology and promote infection. Typically, the sequences of effectors provide little functional information and further targeted experimentation is required. Here, we utilised a structure/function approach to study SnTox3, an effector from the necrotrophic fungal pathogen *Parastagonospora nodorum*, which causes cell death in wheat-lines carrying the sensitivity gene *Snn3*.
- We developed a workflow for the production of SnTox3 in a heterologous host that enabled crystal structure determination. We show this approach can be successfully applied to effectors from other pathogenic fungi. Complementing this, an *in-silico* study uncovered the prevalence of an expanded subclass of effectors from fungi.
- The β-barrel fold of SnTox3 is a novel fold among fungal effectors. We demonstrate that SnTox3 is a pre-pro-protein and that the protease Kex2 removes the pro-domain. Our *in-silico* studies suggest that Kex2-processed pro-domain (designated here as K2PP) effectors are common in fungi, and we demonstrate this experimentally for effectors from *Fusarium oxysporum* f sp. *lycopersici*.
- We propose that K2PP effectors are highly prevalent among fungal effectors. The identification and classification of K2PP effectors has broad implications for the approaches used to study their function in fungal virulence.

## Introduction

*Parastagonospora nodorum* is the causal agent of the wheat disease septoria nodorum blotch (SNB) and is responsible for significant yield losses globally (Murray & Brennan, 2009; Crook *et al.*, 2012; Figueroa *et al.*, 2018). *P. nodorum* is a necrotrophic pathogen that thrives on dead or dying host tissue to cause disease and to reproduce. While initially believed to utilise a suite of cell-wall degrading and lytic enzymes to cause disease, it is now well established that *P. nodorum* secretes a number of proteinaceous effectors (also known as necrotrophic effectors/NEs, or host-selective toxins) (Oliver *et al.*, 2012; McDonald & Solomon, 2018). These effectors promote disease through recognition by corresponding susceptibility gene products present in wheat leading to a programmed cell death response (Oliver *et al.*, 2012). This process is termed effector-triggered susceptibility, and is considered the inverse of the interaction that occurs between biotrophic plant pathogens and their hosts, where recognition of effectors by a corresponding resistance gene product leads to localised programmed cell death in infected cells (Jones & Dangl, 2006). This is an effective mechanism against biotrophic pathogens, which derive their nutrients from living cells and tissues; however, for necrotrophic pathogens cell death is advantageous.

The genetic basis of the *P.* nodorum-wheat interaction has been relatively well described, with a total of nine effector-susceptibility gene interactions identified (Liu *et al.*, 2004; Sarma *et al.*, 2005; Friesen *et al.*, 2006; Friesen *et al.*, 2007; Abeysekara *et al.*, 2009; Liu *et al.*, 2009; Friesen *et al.*, 2012; Liu *et al.*, 2012; Gao *et al.*, 2015; Shi *et al.*, 2015). To date, three effectors have been cloned from *P. nodorum*, SnToxA, SnTox1 and SnTox3; they can induce necrosis, even in the absence of the pathogen, in wheat lines that carry *Tsn1, Snn1* and *Snn3*, respectively (Ballance *et al.*, 1989; Ciuffetti *et al.*, 1997; Friesen *et al.*, 2006; Liu *et al.*, 2009; Faris *et al.*, 2010; Liu *et al.*, 2012). While the identity of Snn3 remains unknown, Tsn1 and Snn1 have been cloned and encode proteins similar to those involved in mediating resistance responses to biotrophic/hemibiotrophic plant pathogens. *Tsn1* encodes a nucleotide-binding oligomerisation domain-like receptor (NLR) with an N-terminal serine/threonine protein kinase domain (Faris *et al.*, 2010), and Snn1 is a member of the wall-associated kinase (WAK) family (Shi *et al.*, 2016). It is thought that *P. nodorum* has acquired the ability to hijack typical defence receptors and downstream pathways involved in resistance against biotrophic/hemibiotrophic pathogens to support its lifestyle (Faris *et al.*, 2010; Shi *et al.*, 2016). Despite this similarity, the molecular mechanisms that lead to effector triggered susceptibility and the molecular functions of necrotrophic effectors are largely unknown.

To further our understanding, we seek to determine the function of SnTox3 in *P. nodorum* pathogenesis. *SnTox3* encodes a 230 amino acid (25.3 kDa) protein with the first 20 amino acids at the N-terminus constituting a signal peptide (Liu *et al.*, 2009). Initial isolation of SnTox3 from culture filtrates identified a ~18 kDa protein in which residues 21-72 could not be detected by tryptic digest mass spectrometry (Liu *et al.*, 2009). On this basis, it was hypothesised that SnTox3, contained a pro-domain that is processed during maturation of the protein prior to secretion. Mature SnTox3 contains six cysteine residues that form three disulfide bonds. At least one of these disulfide bonds is required for activity as dithiothreitol (DTT) treatment prevents SnTox3-induced necrosis in Snn3-containing wheat lines (Liu *et al.*, 2009; Zhang *et al.*, 2017). SnTox3 does not share sequence identity or conserved motifs with any known proteins, and as a result determining its biochemical function has been challenging. However, recent work has identified a direct interaction between SnTox3 and defence-related pathogenesis-related-1 (PR1) proteins from wheat, although the molecular mechanisms underpinning the interaction are yet to be elucidated (Breen *et al.*, 2016).

To gain insight into the function of SnTox3 we determined the three-dimensional structure using X-ray crystallography to a resolution of 1.35 Å, revealing a novel protein fold among fungal effectors. Consistent with previous reports, we confirm that SnTox3 is secreted from *P. nodorum* without the putative N-terminal pro-domain; however, our biochemical studies highlight the importance of this region in SnTox3 protein folding. We demonstrate that specific cleavage of the pro-domain can be achieved *in vitro* using Kex2 protease, and that the removal of the pro-domain dramatically increases SnTox3-induced necrosis in *Snn3-*containing wheat. Kex2 cleavage of pro-domains is not unique to SnTox3 and we demonstrate that Kex2 removes the pro-domain *in vitro* from SnToxA and several of Secreted in Xylem (SIX) effectors from *Fusarium oxysporum* f. sp. *lycopersici* (Fol). Using an *in-silico* approach, we predicted the prevalence of Kex2-processed pro-domain (K2PP) effector proteins, which reveals that a number of effectors from economically important fungal pathogens are putative K2PP effector proteins. Collectively, our findings have broad implications for biochemical and functional studies of many fungal effectors.

## Materials and Methods

### Plant material and fungal and bacterial strains

*Snn3*-containing wheat (*Triticum aestivum* genotype Corack) was grown in a controlled environment chamber with a 16 h day at 20°C and 8 h night at 12°C cycle, and light intensity of 250 μM m^−2^ s^−1^ with 85% relative humidity. *P. nodorum* SN15 was grown on V8-PDA plates and incubated at 22°C under 12 h light cycles for 14 days. Following this, mycelium was harvested and grown at 22°C in liquid Fries 3 medium for 3 days with a 12 h light cycle and constant shaking at 140 RPM. For recombinant expression, SHuffle® T7 Express lysY competent *E. coli* (NEB, C3030J) were cultured at 30°C in Terrific Broth media with appropriate antibiotics for plasmid selection.

### Vector construction

The five effectors used in this study, SnTox3 and SnToxA from *P. nodorum*, and SIX1, SIX4 and SIX6 from *Fusarium oxysporum* f. sp. *lycopersici*, including their putative pro-domains, were codon-optimised for expression in *E. coli* (SnTox3^21-230^, SnToxA^17-178^, FolSIX1^22-284^, FolSIX4^18-242^ and FolSIX6^17-225^) and were introduced into either the pET His6 Sumo TEV LIC cloning vector (2S-T; Addgene #29711) or the modified, Golden Gate-compatible, pOPIN expression vector. For the pET His6 Sumo TEV LIC cloning vector, the resulting constructs contained an N-terminal 6xHis-tag-small ubiquitin modifier (SUMO) fusion followed by a *Tobacco etch virus* (TEV) protease site, and for pOPIN vectors the final constructs contained either an N-terminal 6xHis-tag, 6xHis-tag-SUMO or 6xHis-tag-protein GB1 domain (GB1) followed by a 3C protease cleavage site. For expression studies to determine the importance of pro-domains for effector folding, the effectors excluding their pro-domains (SnTox3^73-230^, SnToxA^61-178^, FolSIX1^96-284^, FolSIX4^59-242^, FolSIX6^62-225^) were cloned into the modified pOPIN expression vector to include a 6xHis-tag-GB1. The Golden Gate digestion/ligation reactions and cycling were carried out as described by (Iverson *et al.*, 2016). All primers and gBlocks were purchased from Integrated DNA Technologies. The integrity of all plasmids was confirmed using Sanger sequencing. For a full list of primers and constructs used in this study see Table S1 and S2.

### Heterologous expression in *E. coli*; protein production and purification

The effectors SnTox3, SnToxA, FolSIX1, FolSIX4 and FolSIX6 were produced in *E. coli* SHuffle®. Bacterial cultures were grown in Terrific Broth media at 30°C with shaking at 225 RPM until OD600 was 0.6-0.8. At this point, the temperature was lowered to 16°C and the cultures induced with isopropyl-1-thio-β-D-galactopyranoside (IPTG) to a final concentration of 500 μM and incubated for a further 16 h. Following centrifugation, cell pellets were resuspended in 50 mM HEPES pH 8.0, 300 mM NaCl, 10 % (v/v) glycerol, 1 mM PMSF and 1 μg/mL DNase. Cells were lysed using an Avestin Emulsifex C5 at ~500-1000 psi. Proteins were purified from the clarified lysate by immobilised metal affinity chromatography (IMAC) on a 5 mL Ni^2+^ His-Trap crude FF column. Fractions containing the protein of interest, as determined by SDS-PAGE analysis, were incubated overnight at 4°C with 6xHis-human rhinovirus 3C protease or 6xHis-TEV protease (~50-100 μg) to cleave the N-terminal fusions. The protein of interest was separated from any uncleaved protein, the fusion tag, and protease by IMAC and purified further by size-exclusion chromatography using either a HiLoad 16/600 Superdex 75 PG column or HiLoad 26/600 Superdex 75 PG column (GE Healthcare) equilibrated with 20 mM HEPES pH 7.5 and 150 mM NaCl. Proteins were concentrated using either a 3 kDa or 10 kDa molecular weight cut-off concentrator (MWCO) (Amicon) and snap-frozen in aliquots using liquid nitrogen prior to storage at −80°C for further use. For SnTox3 crystallisation experiments, an additional anion exchange (AIEX) step was performed prior to SEC by passing the protein over a 1 mL AIEX column (GE Healthcare). SnTox3 protein, following IMAC, was dialysed into a buffer containing 20 mM Tris pH 7 and 100 mM NaCl, prior to passing it over the AIEX column. Under the conditions tested, correctly folded SnTox3 did not bind, however misfolded SnTox3 and contaminants did.

### Kex2 processing of recombinant fungal effector proteins

Recombinant SnTox3^73-230^, SnToxA^17-178^, FolSIX1^22-284^, FolSIX4^18-242^, and FolSIX6^17-225^ were cleaved, to remove their putative pro-domains, using recombinant *Saccharomyces cerevisiae* Kex2 protease (Abcam ab96554) at a 1:200 ratio at room temperature for 48 h. The cleaved protein was purified further using a Superdex Increase 75 10/300 (GE Healthcare Life Sciences), pre-equilibrated with 20 mM Hepes pH 7.5 and 150 mM NaCl.

### Crystallisation, diffraction data collection and crystal structure determination

Initial screening to determine crystallisation conditions for both SnTox3^21-230^ and SnTox3^Kex2^ (Kex2-cleaved SnTox3^21-230^) was performed in 96-well plates (LabTech) at 20°C using the hanging-drop vapour-diffusion method and commercially available sparse matrix screens. For screening, 200 nL drops, which consisted of 100 nL protein solution and 100 nL reservoir solution, were prepared on hanging-drop seals (TTP4150-5100), using a Mosquito robot (TTP LabTech, UK) and equilibrated against 100 μL reservoir solution. The drops were monitored and imaged using the Rock Imager system (Formulatrix, USA) over a period of 21 days. Crystals with the best morphology for SnTox3^21-230^ were observed in 20% w/v PEG 3350 and 0.2 M sodium acetate trihydrate pH 4.5 (ShotGun A3), and 0.1 M HEPES pH 7.5 and 20% w/v PEG 10000 (ShotGun A6). For SnTox3^Kex2^, crystals grew in 0.2 M magnesium chloride hexahydrate, 0.1 M Tris pH 8.0 and 20% w/v PEG 6000 (PACT D10), and 0.2 M sodium nitrate, 0.1 M Bis-Tris propane pH 6.5 and 20% w/v PEG3350 (PACT F5). Crystal optimisation was carried out in 24-well hanging-drop vapour diffusion plate format and involved altering both pH and precipitant concentrations. The final optimised crystallisation conditions for SnTox3^Kex2^ were 0.2 M sodium nitrate, 0.1 M Bis-Tris propane pH 6.5, 18% w/v PEG3350 and 0.2 M magnesium chloride hexahydrate, 0.1 M Tris pH 7.6 and 18-20% w/v PEG 6000.

Before x-ray data collection, crystals were transferred into a cryoprotectant solution containing the reservoir solution with 20% glycerol. SnTox3^Kex2^ crystals were soaked for ~60 s in reservoir solution supplemented with 1 M sodium bromide. Datasets of native and bromide-soaked crystals were collected on the MX2 beamline at the Australian Synchrotron (Table S3). The datasets were processed in XDS (Kabsch, 2010) and scaled using Aimless (Kabsch, 2010; Evans & Murshudov, 2013) in the CCP4 suite (Winn *et al.*, 2011). For bromide-based SAD phasing, the CRANK2 pipeline was used (Skubák & Pannu, 2013) in the CCP4 suite. The model was then refined using phenix.refine in the PHENIX package (Afonine *et al.*, 2012), and iterative model building between refinement rounds was carried out in Coot (Emsley *et al.*, 2010). This model was then used as a template for molecular replacement, using a native SnTox3^Kex2^ dataset. Automatic model building was carried out with AutoBuild (Terwilliger *et al.*, 2008), and the resulting model was refined using phenix.refine in the PHENIX package (Adams *et al.*, 2010), and iterative model building between refinement rounds was carried out in Coot (Emsley *et al.*, 2010). Structure validation was carried out using the MolProbity online server (Davis *et al.*, 2004). A structural similarity search was carried out using the Dali server (Holm & Rosenstrom, 2010).

### Mass spectrometry (MS) of intact protein and N-terminal sequencing to identify point of cleavage in SnTox^321-230^ crystals

MS was carried out using intact SnTox3^21-230^ protein both prior to and after crystallisation, to determine the size of the cleaved protein. The dissolved crystals were buffer-exchanged using a 10 kDa MWCO concentrator (Amicon) to remove the PEG 6000 present in the crystallisation buffer. The samples were then run on an Orbitrap Elite™ (Thermo) mass spectrometer and Dionex UltiMate™ 3000 nano LC system (Thermo). Each sample was first desalted on a PepMap™ 300 C4 pre-column using buffer A (30 μL/min) for 5 min and separated on an Acclaim PepMap 300 (75 um x 150 mm) at a flow rate of 300 nL/min. A gradient of 10-90% buffer B over 5 minutes was used, where buffer A was 0.1% formic acid (FA) in water and buffer B was 80% acetonitrile/ 0.1% FA. The eluted protein was directly analysed on an Orbitrap Elite™ mass spectrometer interfaced with a NanoFlex source. MS was operated in positive ion mode using the Orbitrap analyser set at 60,000 resolution. Source parameters included an ion spray voltage of 2 kV, temperature at 275°C, SID = 30 V, S-lens = 70 V, summed microscans = 10 and FT vacuum = 0.1. MS analysis was performed across 600-2000 m/z. The data was deconvoluted using Thermo Protein Deconvolution™ software across m/z 600-2000, S/N=1, and minimum number of charges set to 3. Deconvoluted data are reported as uncharged monoisotopic masses.

N-terminal sequencing was carried out by the Australian Proteome Analysis Facility (APAF). Approximately 10 μg of SnTox3^21-230^, in a solution containing 10 mM HEPES pH 7.0, 150 mM NaCl, 0.1 M bicine pH 8.0 and 12% PEG 6000, was dissolved in 100 μL of 0.1% trifluoroacetic acid (TFA) and 50 μL of this solution desalted on a ProSorb PVDF filter cartridge (Life Technologies) with three 0.1 % TFA washes. The sample on the PVDF membrane was subjected to 7 residue cycles of Edman N-terminal sequencing using an Applied Biosystems Procise® 494 protein sequencing system.

### Wheat protein-mediated phenotyping assay

SnTox3^21-230^ and SnTox3^Kex2^, at concentrations of 0.1 μM, 0.5 μM, and 1 μM, were syringe-infiltrated into the second leaf of two-week old Corack plants (*Snn3*-containing). Kex2 at a concentration consistent with the highest concentration used for cleavage was used as a control. After 3 days, the leaves were harvested and imaged.

### Western blot analysis of SnTox3 from culture filtrates

For western blot analysis rabbit polyclonal SnTox3 antibodies were generated against recombinant SnTox3^21-230^ protein at the Walter & Eliza Hall Institute of Medical Research (WEHI). Culture filtrates of *P. nodorum* SN15 and a SnTox3KO strain (Liu *et al.*, 2009) were centrifuged at 5000 x *g* for 15 min at 4°C and resolved by SDS-PAGE prior to transfer to a PVDF membrane (BioRad, Hercules, CA, USA). The membrane was then incubated with SnTox3 primary antibodies, which were detected by anti-rabbit IgG HRP from goat (Sigma‐Aldrich). Immunolabelled protein bands were detected on the immunoblot using ECL substrate (BioRad) and visualised by ImageQuant4000 (GE Healthcare).

### Protein lipid overlay assay

Commercial membrane strips, spotted with 100 pmol of various biologically important lipids found in cell membranes (Membrane Lipid Strips, Echelon Biosciences) were used to detect if SnTox3^Kex2^ could bind lipids. SnTox3^Kex2^ was spotted onto the membrane to act as a positive control for antibody detection. Once dried, the membrane was blocked with PBS-T + 1% skim milk powder (blocking buffer). All steps were carried out with agitation at room temperature for 1 h. Then 0.5 μg/mL SnTox3^Kex2^ in blocking buffer was incubated for 1 h, followed by three wash steps using PBS-T for 10 min each. Anti-SnTox3 antibody in blocking buffer (1:1000 dilution) was added and incubated for 1 h, followed by three wash steps as described previously. The bound antibody was detected with an anti-rabbit IgG HRP from goat (1:2000 dilution) in 10 mL of PBS-T, followed by a wash step, as described. Protein binding was detected by using ECL substrate (BioRad) and visualised by ImageQuant4000 (GE Healthcare, Silverwater, TX, USA).

### Bioinformatics search for effectors with Kex2-processed pro-domains

A python script was used to search for the occurrence of Kex2-processed pro-domains (K2PP) in secreted fungal effectors (https://github.com/JanaSperschneider/Publications_Code/tree/master/2020_04_Tox3_LxxR_Paper). First, SignalP 3.0 (Bendtsen *et al.*, 2004) was run to determine the predicted signal peptide cleavage site. We then searched for occurrences of LxxR, KR or RR motifs in the predicted secreted protein. For all motif positions, we analysed the N-terminal sequence before the motif (excluding the predicted signal peptide) and the C-terminal sequence after the motif. If the N-terminal sequence was longer than four amino acids and the C-terminal sequence occupied more than half of the mature secreted protein, it was analysed further. We analysed the percentages of amino acids that are associated with disorder (K, E, N, S, P, G, R, D, Q, M) and those that are associated with order (W, Y, F, I, C, L, V, H) (Weathers *et al.*, 2004) in the N-terminal sequence and in the C-terminal sequence. If at least two-thirds of amino acids (aas) in the N-terminal sequence are disorder-promoting (disorder-promoting aas/(disorder-promoting aas + order-promoting aas)) and if the proportion of disorder-promoting amino acids in the N-terminal sequences is higher than in the C-terminal sequence, the secreted protein was labelled as having a predicted disordered region with a Kex2 protease cleavage site. The list of fungal effectors were taken from the EffectorP 2.0 publication (Sperschneider *et al.*, 2018).

### Agro-infiltration of SnTox3 in *Nicotiana benthamiana*

The SnTox3-SP:GFP construct was generated by recombining pDONR201-PR1ΔSPSnTox3 (Breen *et al.*, 2016) into the plant expression vector pB7FWG2.0 using a Gateway LR reaction. The pB7WGF2.0_empty vector (EV:GFP) was used as a negative control. After sequence confirmation, the constructs were expressed in *Nicotiana benthamiana* using agro-infiltration, as described previously.

## Results

### The effector SnTox3 is synthesised as a pre-pro-protein

Previously, we reported a protocol for producing several functionally active fungal effectors, including SnTox3^21-230^ (pro-domain containing) using the *E. coli* strain SHuffle® (Zhang *et al.*, 2017). However, the overall yields for SnTox3 were low and not amenable for structural studies. To address this problem, we tested the ability of two N-terminal cleavable fusion partners, small ubiquitin-like modifier (SUMO) and protein GB1 domain (GB1), to enhance solubility and improve SnTox3 yields compared to a 6xHis tag alone. The addition of both 6xHis-SUMO and 6xHis-GB1 improved overall yields, with comparable purity, and the GB1 fusion resulted in the highest yields (Fig. **S1**, Table **S4**). The improvements made to SnTox3 protein production enabled us to procced with crystallisation studies and thin, needle-like crystals were obtained for SnTox3 (data not shown). However, these crystals diffracted poorly (>9 Å resolution), despite extensive optimisation, preventing structure determination.

We suspected additional processing of SnTox3 may be influencing crystal formation and quality. SDS-PAGE analysis of the protein from the crystallisation drops revealed that the majority of SnTox3 was ~6 kDa smaller than expected (Fig. **1a**), suggesting that proteolytic cleavage was occurring post purification. Characterisation of the protein in the crystallisation drop using intact MS identified a species with a monoisotopic mass of ~18 kDa (Fig. **S2**). Further analysis using N-terminal sequencing demonstrated that this ~18 kDa protein began at residue Tyrosine 73 (Fig. **1b**), which corresponds to the sequence immediately after the putative Kex2 protease recognition site identified previously by Liu *et al.* (2009).

**Figure 1:**
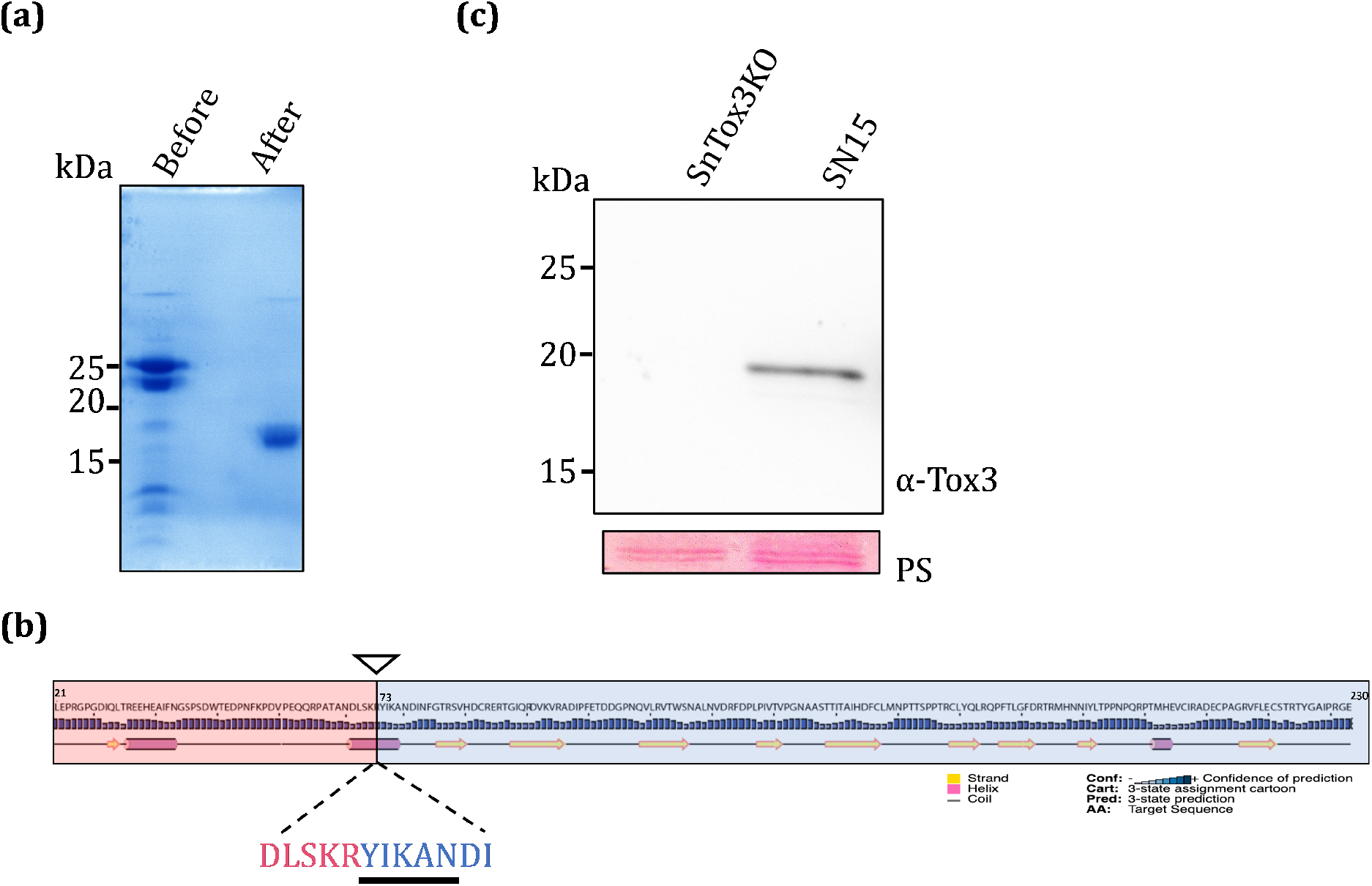
SnTox3 is secreted as a pre-pro-protein from *P. nodorum*. **(a)** Coomassie-stained SDS-PAGE gel showing recombinant SnTox3^21-230^ prior to crystallisation and after crystal formation shows a difference in size of ~5-10 kDa. **(b)** Schematic diagram of SnTox3^21-230^, highlighting secondary structure predictions as determined by PSIPRED (Jones, 1999; Buchan & Jones, 2019) with confidence levels shown as a bar graph. The putative pro-domain (residues 21-72; 5.9 kDa) is shown in red, and the mature domain (73-230; 17.9 kDa) is shown in blue. The black arrow indicates the putative Kex2 cleavage site, and the first five residues detected by N-terminal sequencing of the protein from the crystallisation drop are enlarged and underlined in black. **(c)** Top panel: culture filtrates of *P. nodorum* SN15 and SnTox3KO strains were analysed by western blot and probed with SnTox3 primary antibodies, which were detected by goat anti-rabbit IgG HRP. Bottom panel: Ponceau staining (PS) to show protein loading. Numbers on the left of gel images indicate sizes of molecular weight markers.

We subsequently sought to determine what form of SnTox3 is produced by the fungus. Antibodies, generated against purified SnTox3 protein, were used in western blot analysis of culture filtrates of *P. nodorum* isolate SN15 and a SnTox3 knockout variant (Fig. **1c**). This analysis demonstrated that the secreted form of SnTox3 corresponds to an 18 kDa protein, indicating that the pro-domain is removed to produce the mature protein found in the culture filtrate, which is consistent with removal of the signal peptide and pro-domain prior to secretion.

### The SnTox3 pro-domain is necessary for SnTox3 heterologous production but reduces necrosis-inducing activity

In light of these findings, we sought to produce recombinant SnTox3 without the pro-domain (SnTox3^73-230^). Strikingly, we observed a substantial reduction (~10X less) in the amount of soluble SnTox3^73-230^ compared to SnTox3 with the pro-domain (SnTox3^21-230^), regardless of the use of a GB1 fusion (Fig. **S3a**). These data suggest that the pro-domain is essential for the production of correctly folded SnTox3 in *E. coli*. To overcome this obstacle, we pursued removal of the pro-domain using Kex2 to mimic the natural processing and maturation of SnTox3. Kex2 is an important endogenous protease conserved across fungi (Wickner, 1974; Newport & Agabian, 1997; Bader *et al.*, 2008; Jacob-Wilk *et al.*, 2009), which cleaves pro-proteins in a site-specific manner. The cleavage motif for Kex2 is typically described as a dibasic motif, with a preference for Arg at position P1 (where the order is P4‐P3‐P2‐P1‐cleavage) and a basic residue (typically Lys or Arg) at position P2 (Bevan *et al.*, 1998; Bader *et al.*, 2008). Additional *in vivo* characterisation of this cleavage motif in *S. cerevisiae* has suggested that specificity also occurs at P4, with a particular preference for Leu or other aliphatic residues (Rockwell & Fuller, 1998; Li *et al.*, 2017). In SnTox3, a LSKR motif is localised at the pro-domain/mature protein junction (Fig. **1c**, Table **S4**). To test if Kex2 can remove the pro-domain from SnTox3, we incubated Kex2 protease with purified SnTox3^21-230^ and demonstrated the selective removal of the N-terminal pro-domain (Fig. **2a**), further implicating the role of Kex2 in effector maturation. We subsequently defined SnTox3 as a Kex2-processed pro-domain (K2PP) effector.

**Figure 2:**
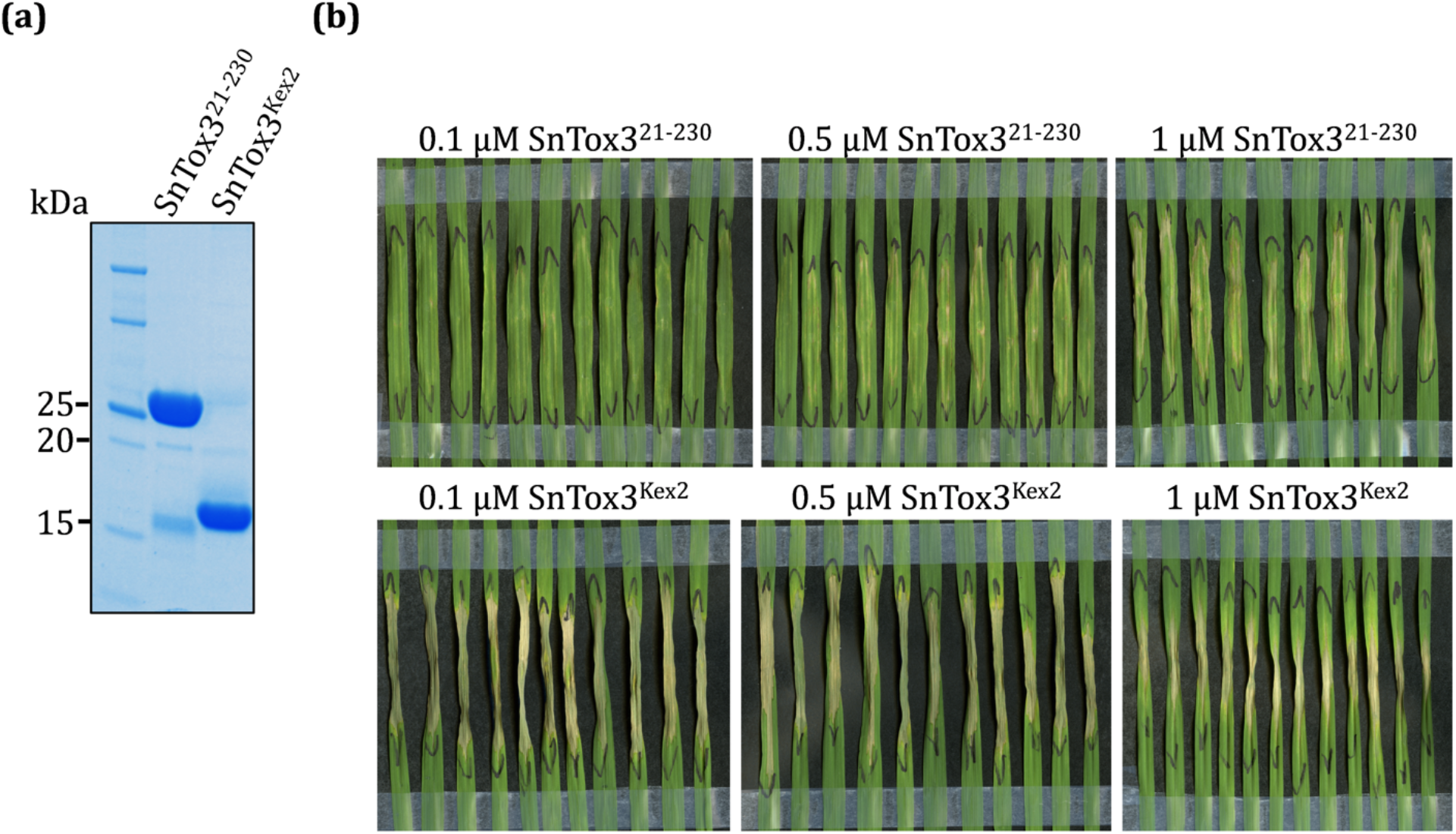
Removal of the pro-domain from recombinant SnTox3 increases necrosis-causing activity. **(a)** Coomassie-stained SDS-PAGE gel showing *in vitro* Kex2-mediated cleavage of SnTox3^21-230^ using a 1/200 dilution (of SnTox3^21-230^: Kex2 (Abcam ab96554)). The first lane shows molecular weight markers, with sizes indicated by numbers on the left-hand side. Lane 2 contains recombinant SnTox3^21-230^ and lane 3 is SnTox3^21-230^ following cleavage by Kex2 (Tox3^Kex2^). **(b)** Necrosis caused by 0.1, 0.5 and 1 μM SnTox3^21-230^ (top panel) or SnTox3^Kex2^ (bottom panel) following infiltration into the second leaf of Corack (*Snn3*-containing) wheat leaves. The black lines indicate the boundary of the infiltration zones. Leaves were harvested and photographed at 3 days post-infiltration (dpi).

To understand the effect on the necrosis-inducing activity of SnTox3 following pro-domain removal, we infiltrated recombinant SnTox3^21-230^ and SnTox3^Kex2^ (pro-domain removed with Kex2), at concentrations of 0.1, 0.5, and 1 μM, into the 2^nd^ leaf of 2-week-old Corack (*Snn3*-containing) seedlings (Fig. **2b**). After 3 days, the leaves infiltrated with SnTox3^Kex2^ showed more advanced and severe signs of necrosis across the tested concentrations when compared to full-length SnTox3 (Fig. **2b**). Importantly, Kex2 alone did not induce cell death (Fig. **S4**). These data demonstrate that inclusion of the pro-domain has an inhibitory effect on the necrosis-causing activity of SnTox3, which likely explains why this region is removed during maturation and secretion of the protein.

### The crystal structure of SnTox3 reveals a β-barrel fold

With the role of the pro-domain defined for SnTox3, we decided to use SnTox3^Kex2^ for crystallisation. This approach was successful and enabled us to determine the crystal structure of SnTox3^Kex2^ to a resolution of 1.35 Å, using a bromide ion-based single-wavelength anomalous diffraction (SAD) approach (Fig. **3a**, Table **S3**). Overall, mature SnTox3 contains ten β-strands (β1-β10), where eight of the β-strands are connected in an antiparallel up-and-down topology by loops of various lengths and result in a β-barrel, linked together by three disulfide bonds. The only region not bound by disulfide bonds back to the barrel includes the β-strands β3 and β4, encompassing residues 113 to 137, which adopt a β-hairpin-like structure (Fig. **3a**). Interpretable electron density for three disulfide bonds was observed and shows that they all localise to one end of the protein (Fig. **3a**, **3b**). The positioning of the disulfide bonds, within the context of the fold, suggests that these bonds play a role in providing overall stability to the structure, and to specifically anchor the β-strands together. The connectivity of the disulfide bonds differs from that originally predicted for SnTox3 (Liu *et al.*, 2009); disulfide bonds are formed by the pairs C89-C218, C154-C209, and C166-C203, form disulfide bonds (Fig. **3a, 3b**).

**Figure 3:**
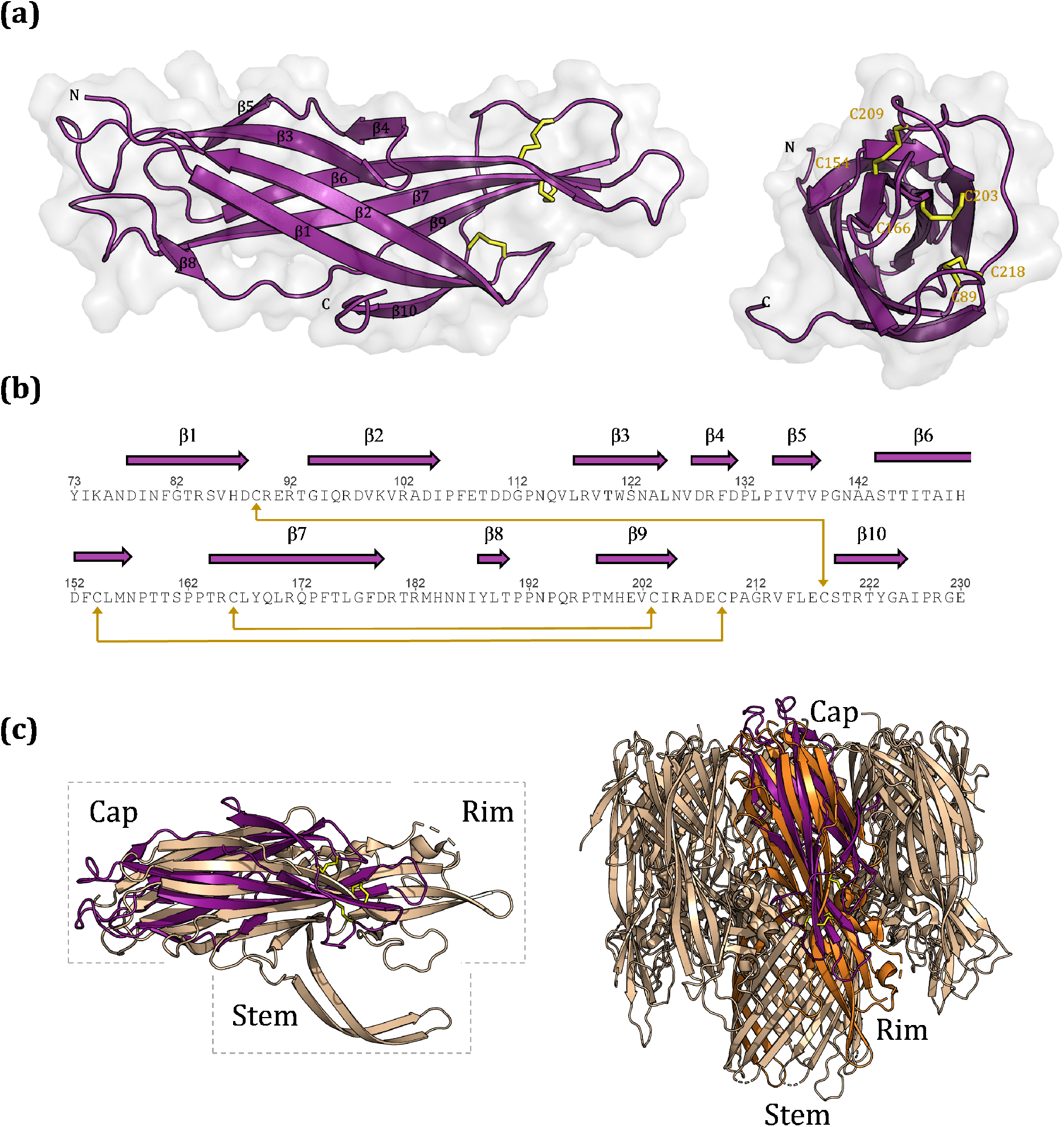
Crystal structure of SnTox3 shows a novel effector fold. **(a)** Ribbon diagram of SnTox3, showing the overall β-barrel fold, with disulfide bonds shown in yellow. Disulfide-bond connectivities are shown on the SnTox3 ribbon diagram, with the residues labelled accordingly. **(b)** The amino-acid sequence of SnTox3, showing secondary structure elements; disulfide bond connectivity is shown with gold lines. **(c)** Structure superimposition of SnTox3 (purple) and LukGH (PDB ID: 4TW1; gold). Left panel: superposition of SnTox3 (purple) with one protomer of LukGH (chain A), showing structural similarity with only the cap domain and not the rim and stem domains, which are required for pore formation. Right panel: superimposition of SnTox3 (purple) with one protomer (orange) in the pore complex of LukGH (gold).

### SnTox3 structure is novel among fungal effectors but shares structural similarity with bacterial pore-forming toxins (PFTs)

To identify whether SnTox3 was structurally similar to other proteins, the structure was compared against all reported structures in the Protein Data Bank utilising the Dali server (Holm & Rosenstrom, 2010). SnTox3 shares only low structural similarities with proteins of known structure and no structural similarity to other fungal effectors (Fig. **S5**). The most similar structures to SnTox3 are the family of bacterial pore-forming toxins (PFTs), originating from various pathogenic bacterial species (Fig. **S5**). In particular, SnTox3 shares similarity to β-PFTs, which form a β-barrel structure that inserts into lipid bilayer to form the pore. The closest structural match was the bi-component toxin LukGH (leukocidin) from *Staphylococcus aureus*. Structure superposition (Fig. **3c**) of SnTox3 and LukGH reveals a root-mean-square deviation (RMSD) of 3.7 Å for 107 structurally equivalent Cα atoms, despite sharing <10% protein sequence identity (Fig. **S5**). The structural similarity between LukGH and SnTox3 is confined to the cap domain of LukGH only, which is the extracellular domain that interacts with adjacent protomers in the pore complex (Menestrina *et al.*, 2003; Parker & Feil, 2005; Badarau *et al.*, 2015) (Fig. **3c**). There are also noticeable differences between SnTox3 and the cap domains. Cap domains in PFTs adopt an overall β-sandwich fold consisting of two, generally six-stranded, antiparallel β-sheets (Menestrina *et al.*, 2003; Parker & Feil, 2005; Badarau *et al.*, 2015), whereas SnTox3 contains ten β-strands that form a β-barrel. In light of the structural differences and the lack of the rim and stem domain, SnTox3 alone would be unable to form a membrane-spanning pore analogous to PFTs. Despite these differences, we tested whether SnTox3 would associate with lipids using a simple lipid overlay assay, as reported for several bacterial PFTs (Savva *et al.*, 2013; Gil *et al.*, 2015). No binding of SnTox3^Kex2^ was observed to any of the membrane lipids tested (Fig. **3d**).

### Predicted Kex2-processed pro-domains are common in fungal effectors

Several reports have implicated Kex2 in the processing of N-terminal regions from other fungal effectors (Jia *et al.*, 2000; Basse *et al.*, 2002; Rep, 2005; Houterman *et al.*, 2007; Simbaqueba *et al.*, 2018). We wished to determine if our approach for protein production of SnTox3 was more broadly applicable to other cysteine-rich fungal effectors. Four effectors, including SIX1, SIX4 and SIX6 from *Fusarium oxysporum* f. sp. *lycopersici* (Fol) and SnToxA were used for further experimentation (Table **S4**). Previously, FolSIX1 and FolSIX4 were suggested to be synthesised as pre-pro-proteins following identification of a putative Kex2-processing site (Rep, 2005; Houterman *et al.*, 2007), and SnToxA is thought to be synthesised as a pre-pro-protein (Ballance *et al.*, 1989; Ciuffetti *et al.*, 1997), although Kex2 was never implicated in SnToxA processing. We produced these effectors using *E. coli* SHuffle, and found that the use of an N-terminal GB1 fusion improved soluble protein yields in all cases, consistent with our data for SnTox3 (Fig. **S6-S9**, Table **S4**). We also found that the putative pro-domain was required to produce all effector proteins in a soluble form using the *E. coli* expression system (Fig. **S3**). We subsequently demonstrated that Kex2 could remove the N-terminal putative pro-domain of all four effectors (Fig. **4a**). For FolSIX4 and FolSIX1, some additional processing was observed after Kex2 treatment, which is consistent with the presence of multiple putative Kex2 motifs (Lys-Arg and Arg-Arg) within these protein sequences (Table **S5**). Collectively, these data are consistent with these proteins representing K2PP effectors and suggest that the pro-domains are essential for producing correctly folded proteins.

**Figure 4:**
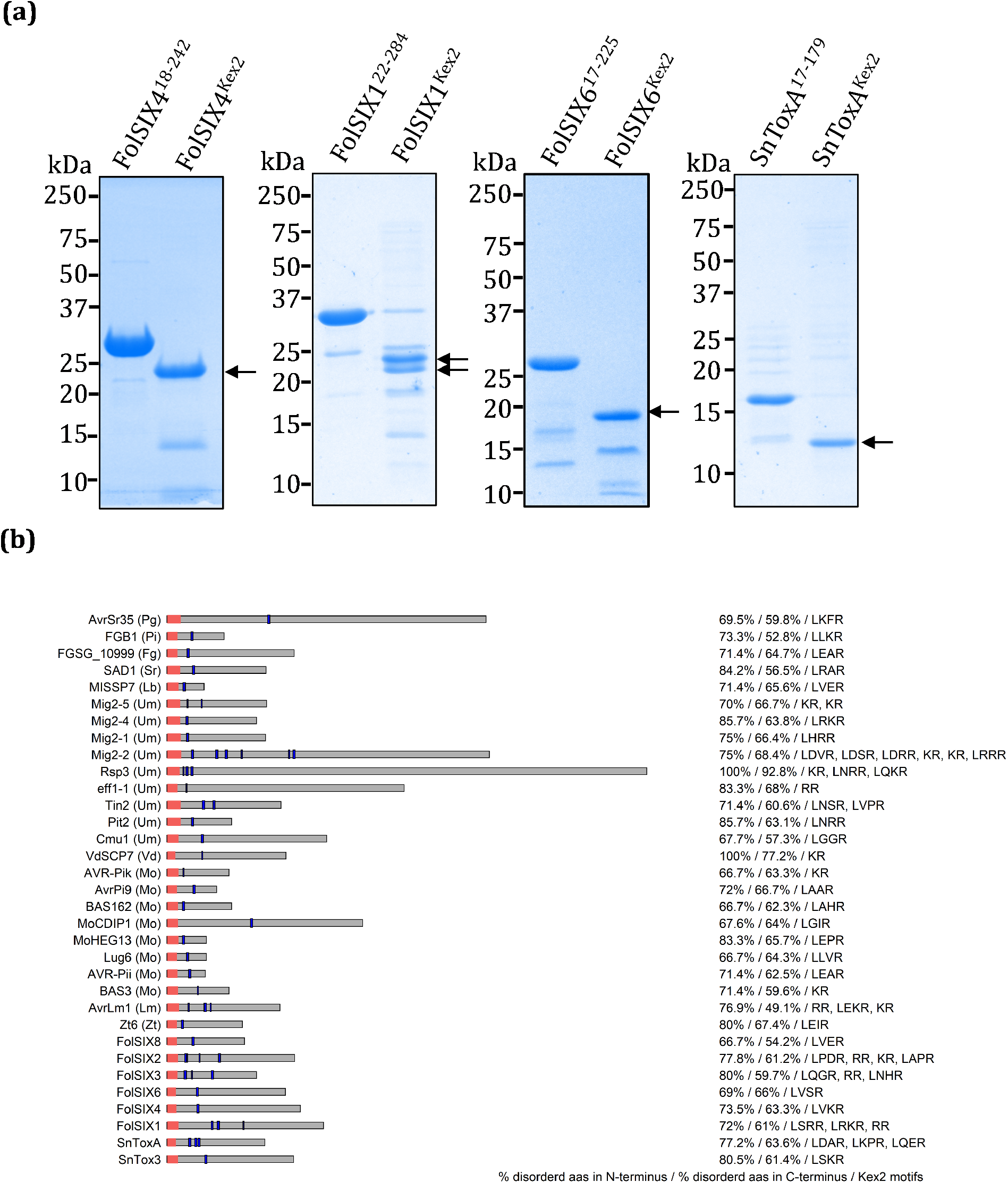
Fungal effectors with predicted Kex2-processed pro-domains. **(a)** Coomassie-stained SDS-PAGE gel showing *in vitro* Kex2-mediated cleavage of FolSIX6^17-225^, FolSIX4^18-242^, FolSIX1^22-284^, and ToxA^17-178^. Numbers on the left-hand side indicate the sizes of molecular weight markers **(b)** The 33 of 120 fungal effectors that are predicted to carry LxxR, KR or RR motifs that are preceded by disordered domains using an *in-silico* approach. Effectors are shown with red boxes indicating the signal peptide region and blue boxes indicating the LxxR, KR or RR motifs. The fungal species from which the effector originates is indicated; *Puccinia graminis* (Pg), *Piriformospora indica* (Pi), *Fusarium graminearum* (Fg), *Sporisorium reilianum* (Sr), *Laccaria bicolor* (Lb), *Ustilago maydis* (Um), *Verticillium dahlia* (Vd), *Magnaporthe oryzae* (Mo), *Leptosphaeria maculans* (Lm), *Zymoseptoria tritici* (Zt), *Fusarium oxysporum f. sp. lycopersici* (Fol), and *Parastagonospora nodorum* (Sn). For each effector, the amino acid composition of the N-terminal region (sequence from signal peptide cleavage site to start of first Kex2 motif) and the C-terminal region (end of last Kex2 motif until end of sequence) are analysed. The proportion of disorder-promoting amino acids is shown on the right, along with all putative Kex2 cleavage sites that occur in the first half of the protein sequence.

Based on our *in vitro* studies, we investigated the prevalence of K2PPs in fungal effectors using an *in-silico* approach. We analysed the sequences of FolSIX1, FolSIX4, FolSIX6, SnTox3 and SnToxA to uncover common features. We identified a conserved Leu at P4, and Arg at P1 with varied residues at P3 and P2 (LxxR) at the junction of the putative pro-domains and mature domain. This is consistent with a recent study from yeast that defined a major cleavage pattern for Kex2 protease as being an aliphatic (preferentially a Leu) at P4 and either a Lys or Arg at P2, followed by an Arg at P1 (Li *et al.*, 2017). In addition, pro-domains in the experimentally defined effectors are predicted to be predominantly disordered. As a result, we searched for the presence of LxxR, and the canonical Kex2 dibasic (KR and RR) motifs in 120 fungal effectors taken from Sperschneider *et al.* (2018). Given the prevalence of these motifs in the effector protein sequences we further constrained our search to within the first half of the amino acid sequence (following signal peptide removal) and included a criterion whereby the motif needed to be preceded by a high proportion of disorder-promoting amino acids. Of these effectors, 33 (27.5%) are predicted to have K2PPs (Fig. **4b**), including all effectors previously implicated as being Kex2 cleaved in the literature, except Avr-Pita (Jia *et al.*, 2000; Rep, 2005). In several effectors multiple putative Kex2 motifs were identified to be present (Fig. **4b**, Table **S5**) and in some instances the presence of a potential Kex2 motifs in the mature domain of the protein was observed (see Table S5).

## Discussion

To date, several structures of effector proteins originating from fungal plant pathogens have been determined, and have provided insights into the biochemical functions of these proteins as well as how they are recognised by the plant (Sarma *et al.*, 2005; Guncar *et al.*, 2007; Wang *et al.*, 2007; Sánchez-Vallet *et al.*, 2013; Ve *et al.*, 2013; Nyarko *et al.*, 2014; de Guillen *et al.*, 2015; Maqbool *et al.*, 2015; Ose *et al.*, 2015; Liu *et al.*, 2016; Di *et al.*, 2017; De la Concepcion *et al.*, 2018; Hurlburt *et al.*, 2018; Zhang *et al.*, 2018). An emerging insight from these structures is the existence of conserved folds despite originating from fungi that belong to different taxa and having high levels of sequence diversity (reviewed by (Franceschetti *et al.*, 2017)). An example is the MAX (*Magnaporthe* Avrs and ToxB-like) effector family, which includes representatives from the rice blast pathogen *M. oryzae* (Zhang *et al.*, 2013; de Guillen *et al.*, 2015; Maqbool *et al.*, 2015; Ose *et al.*, 2015) and the wheat pathogen *Pyrenophora tritici-repentis* (Nyarko *et al.*, 2014). Here, we report the first effector structure from *P. nodorum* and show that the β-barrel fold of SnTox3 is novel among fungal effectors, perhaps suggestive of a new structural family. In general, β-barrel folds are known to be associated with high stability and robustness against temperature and pH changes (Koebnik *et al.*, 2000; Tamm *et al.*, 2004). SnTox3 is proposed to function in the apoplast (Liu *et al.*, 2009) and its structure likely plays an important role in maintaining protein stability within this environment. Protein stability is also enhanced by the formation of three disulfide bonds, which link the β-strands together, and in light of the structure, it is not surprising that reduction of these bonds via DTT treatment leads to the abolition of SnTox3-induced cell death (Liu *et al.*, 2009).

Our structural similarity searches revealed that SnTox3 shares the highest structural similarity with the cap domains of bacterial pore-forming toxins (PFTs), despite sharing no identifiable similarity at the protein sequence level. While intriguing, particularly given commonalities in cytotoxic function and pathogen virulence, the lack of important additional domains in SnTox3 demonstrate that the protein alone is unlikely to perforate membranes. At this stage, the structure of SnTox3 has not enabled us to directly infer function. This has been the case with most fungal effector structures published to date (Sarma *et al.*, 2005; Wang *et al.*, 2007; Ve *et al.*, 2013; Nyarko *et al.*, 2014; Blondeau *et al.*, 2015; de Guillen *et al.*, 2015; Maqbool *et al.*, 2015), where additional protein-protein interaction screening and biochemical experiments have been needed to derive function.

Here, we show experimentally that SnTox3 contains a Kex2-processed pro-domain. Kex2 is a highly-conserved serine protease that localises to the late trans-Golgi network (Redding *et al.*, 1991) and a pre-vacuolar compartment in fungi (Blanchette *et al.*, 2004). Kex2 is responsible for the maturation of proteins in a site-specific manner and plays a pivotal role in protein secretion in yeast and fungi. Recently, the prevalence of Kex2-processed repeat proteins in nearly all fungi, including human and plant pathogens, was shown using a genome-wide survey of 250 publicly available fungal secretomes (Le Marquer *et al.*, 2019). Some of the identified peptides corresponded to sexual pheromones, mycotoxins as well as effector proteins, and many of the putative-processed protein had unknown functions (Le Marquer *et al.*, 2019). While Kex2 processing is not a defining feature of fungal effectors, a role for the protease has been implicated in pathogen virulence. Several Kex2 deletion mutants have been studied in human pathogens, including *Candida albicans*, *Aspergillus niger*, and *Cryphonectria parasitica* (Newport & Agabian, 1997; Newport *et al.*, 2003; Punt *et al.*, 2003; Jacob-Wilk *et al.*, 2009). Collectively in these mutants, reduced virulence was observed but they also suffered from other pleotropic effects and morphological changes, highlighting the general role that Kex2 plays in fungal growth and development.

In fungal effectors, Kex2 cleavage of repeat-containing effectors, such as Rep1, Hum3, and Rsp1 from *Ustilago maydis*, is known to play a role in pathogen infection and virulence (Wösten *et al.*, 1996; Müller *et al.*, 2008; Mesarich *et al.*, 2015; Ma *et al.*, 2018). Despite this, to the best of our knowledge, Kex2 processing to remove the N-terminal pro-domain of a fungal effector has not been demonstrated experimentally prior to this report. The current literature infers the role of Kex2 by highlighting the absence of the N-terminal pro-domain region in the mature form of the protein based on mass spectrometry experiments and the association with ‘canonical’ dibasic Kex2 recognition sites (Ballance *et al.*, 1989; Ciuffetti *et al.*, 1997; Jia *et al.*, 2000; Basse *et al.*, 2002; Rep, 2005; Houterman *et al.*, 2007; Simbaqueba *et al.*, 2018). Based on our data, we suggest that K2PP effectors are highly prevalent in fungal effectors and this has broad implications in fungal effector biology.

Pro-domains are found in a diverse range of proteins, and are best known for their roles in controlling activity in proteases and hormones. They have also been implicated in stabilisation and correct folding of these proteins (Baker *et al.*, 1993; Zanin *et al.*, 2017). We found that inclusion of the pro-domain is crucial for producing soluble cysteine-rich effector proteins in *E. coli*. This agrees with early studies involving PtrToxA, whereby the pro-domain was required for protein refolding when produced as an insoluble protein in *E. coli* (Tuori *et al.*, 2000). A number of studies involving effectors from our K2PP effector list (Fig. **4b**) have reported consequences when manipulating the putative pro-domain. For example, deletion of residues 24-60 in the *U. maydis* effector Rsp3 prevented effector secretion and led to protein accumulation within fungal cells (Ma *et al.*, 2018). Similarly, removal of the putative pro-domain from the *Zymoseptoria triti* effector Zt6 led to loss of protein function. It was suggested that this region was involved in protein re-entry into host cells (Kettles *et al.*, 2018); however, it remains plausible that inappropriate trafficking or protein misfolding caused loss of activity. Collectively, the available data suggest that pro-domains are involved in protein folding and trafficking of K2PP effectors, but clearly further experimentation is required.

The removal of the pro-domain (post-folding) was pivotal for obtaining high quality crystals of SnTox3. We also observed that the activity of SnTox3 was hindered if the pro-domain remained intact (Fig. **2b**). This brings to light important considerations for studying K2PP fungal effectors. Kex2 is specific to fungi, and is not present in plants. K2PP effectors are often studied and produced *in-planta* via transient or stable expression without taking account of their Kex2 processing requirements. We found that in *Nicotiana benthamiana*, the pro-domain of SnTox3 is not cleaved during secretion, demonstrating that the protein is not matured *in planta* in the same way as it is within the fungus (Fig. **S10**, Fig. **1c**). Another consideration that should be made is immune-reactive tag positioning (N or C-terminus) and how this could be impacted by the processing of the pro-domain, particularly in fungal expression systems or if Kex2 is utilised *in vitro* for effector maturation. These examples represent but a few crucial considerations when studying K2PP effectors.

## Conclusions

Studies aiming to understand the molecular mechanisms of how plant pathogen effectors modulate host physiology and defence pathways is a major focus in the field of plant-microbe interactions. Here, we report the crystal structure of SnTox3, which has a novel fold among fungal effectors. SnTox3 is a Kex2-processed pro-domain (K2PP) effector, and we demonstrate that K2PPs are present in significant proportion of fungal effectors. Our work with SnTox3 and other pro-domain containing effectors provides a template for the production and study of K2PP effectors in general, which has broad implications for other researchers characterising fungal effectors using both *in vitro* and *in vivo* methods.

## Acknowledgements

This work was supported by the Australian Research Council (ARC; DP160102244 and DP190102526 to B.K., and DP120103558 and DP180102355 P.S.). S.W. was funded by an ARC DECRA (DE160100893) and is supported by the ANU Future Scheme (35665). B.K. was an NHMRC Principal Research Fellow (1110971) and is an ARC Laureate Fellow (FL180100109). P.S. was an ARC Future Fellow (FT110100698). J.S. is funded by an ARC DECRA (DE190100066). M.O. was a recipient of the Australian Government Research Training Program (RTP) Stipend Scholarship. D.Y. was a recipient of the AINSE Honours Scholarship Program. S.R. was a recipient of an Australian Government Research Training Program International Fee Offset Scholarship. We thank Mark Youles in the TSL Synbio team, Adam Bentham, and Mark Banfield for providing the pOPIN golden gate vectors. The MS analysis was carried out at the Mass Spectrometry Facility in the School of Chemistry and Molecular Biosciences, University of Queensland and we thank Peter Josh and Amanda Nouwens for their technical assistance. We acknowledge the use of the University of Queensland Remote Operation Crystallization and X-ray (UQ ROCX) facility at the Centre for Microscopy and Microanalysis and the support from staff, Gordon King and Karl Byriel. We also acknowledge use of the Australian Synchrotron MX facility and thank the staff for their support. Aspects of this research have been facilitated by access to the Australian Proteome Analysis Facility supported under the Australian Government’s National Collaborative Research Infrastructure Strategy (NCRIS). The co-ordinates and structure factors for SnTox3 have been deposited in the PDB with accession number 6WES.

## Author contributions

M.O., D.J., P.S., B.K., and S.W. designed the study; M.O., Y.S., D.Y., B.D., S.R., D.E., J.S and S.W. performed the experiments; all authors analysed the data. M.O. and S.W. wrote the original draft and all authors contributed to writing, reviewing and editing of the manuscript.

