## Supplemental data for "The crystal structure of SnTox3 from the necrotrophic fungus *Parastagonospora nodorum* reveals a unique effector fold and insights into Kex2 protease processing of fungal effectors"

[1]Research School of Biology, The Australian National University, Canberra, ACT 2601, Australia [2]School of Chemistry and Molecular Biosciences, Institute for Molecular Bioscience and Australian Infectious Diseases Research Centre, University of Queensland, Brisbane, Queensland 4072, Australia, [3]Australian Synchrotron, Macromolecular Crystallography, Clayton, Victoria 3168, Australia, [4]Biological Data Science Institute, The Australian National University, Canberra, ACT 2601, Australia

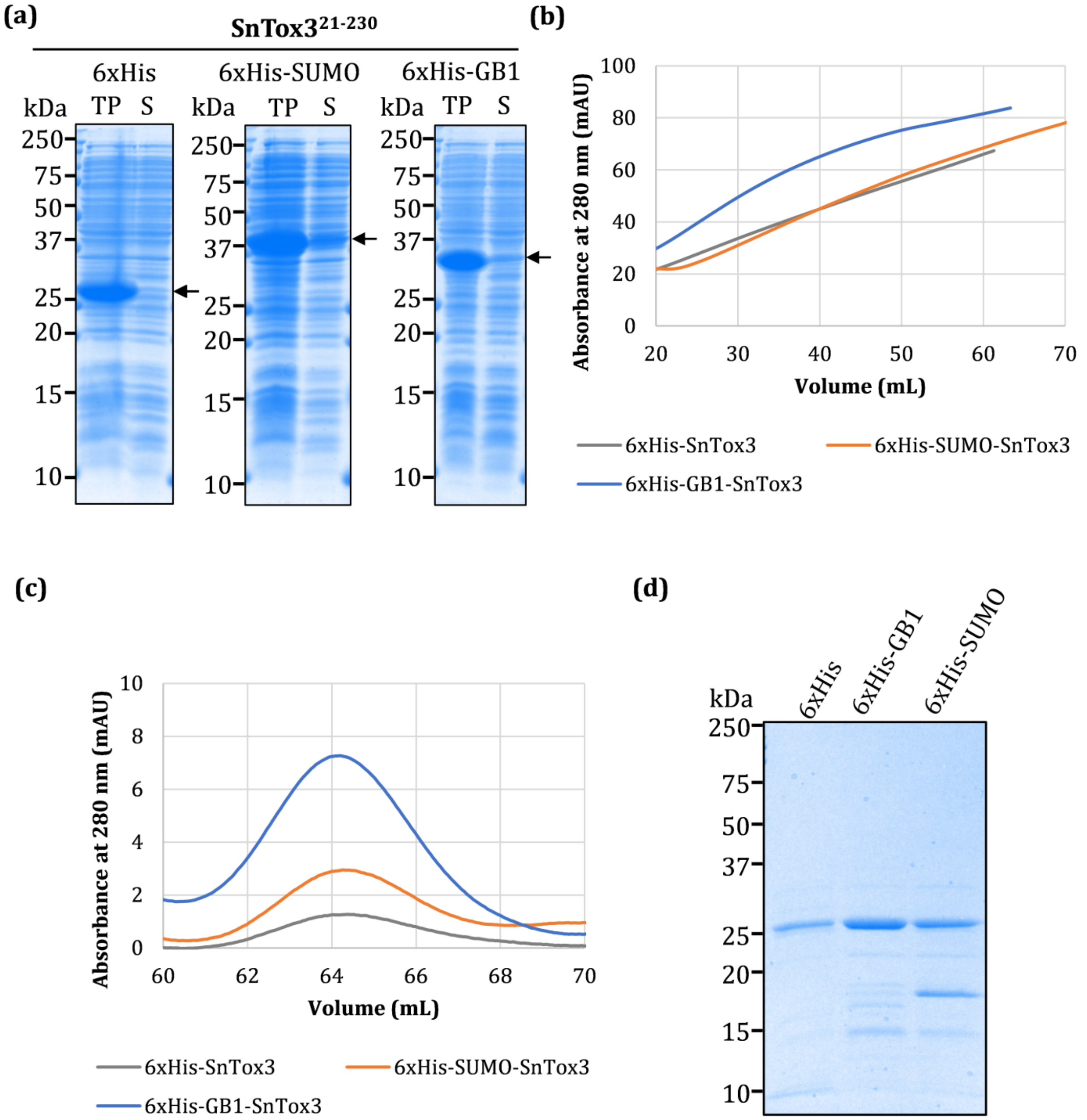


**Fig. S1 Comparison of fusion partners for SnTox3$^{21\text{-}230}$ protein production. (a)** Coomassie-stained SDS-PAGE gel showing total protein (TP) and the soluble fraction (S) for each fusion tag (6xHis-SnTox3, 6xHis-SUMO-SnTox3 and 6xHis-GB1-SnTox3). **(b)** Nickel affinity chromatography profiles for the three SnTox3-fusion protein constructs. **(c)** Size exclusion

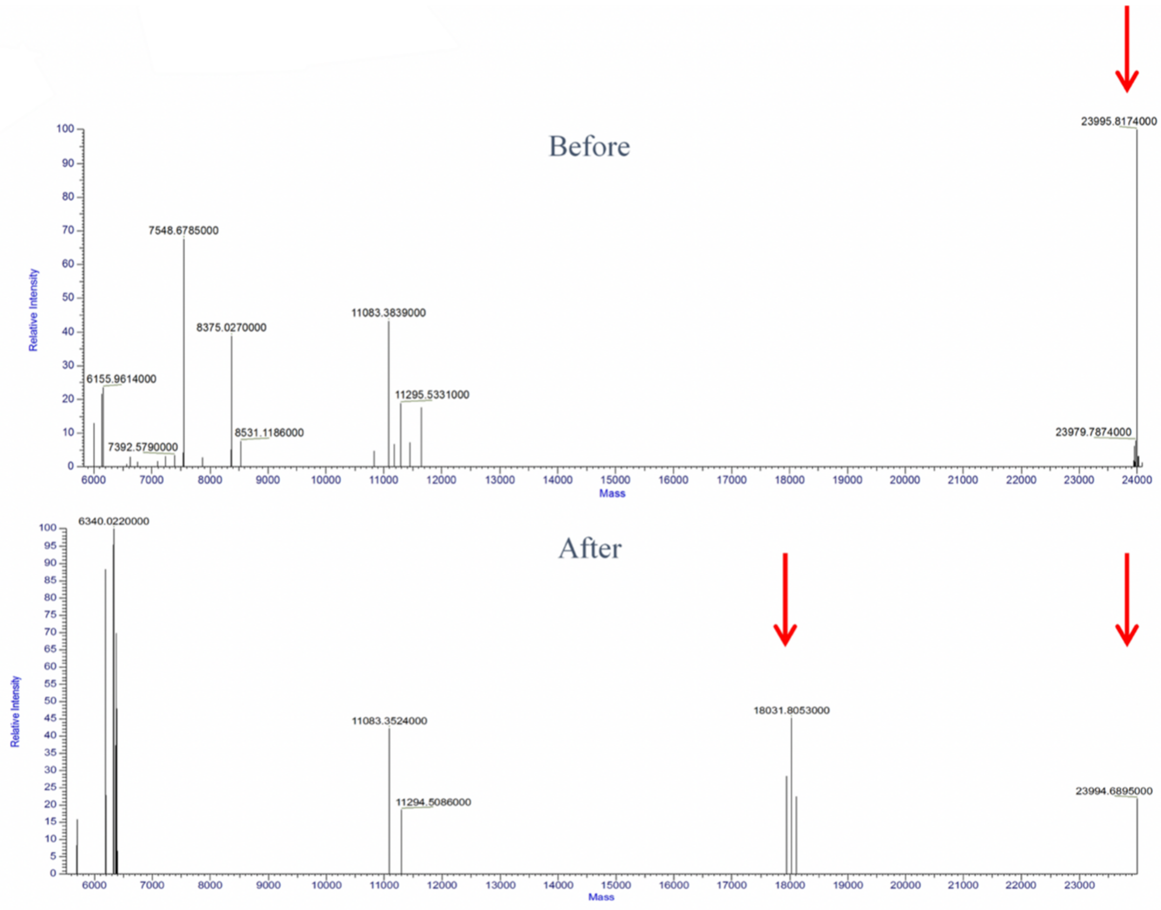


chromatography (SEC) profiles for the three SnTox3-fusion protein constructs, following tag cleavage and removal of uncleaved protein and contaminants by a second nickel affinity chromatography step. **(d)** Coomassie-stained SDS-PAGE gel showing purified SnTox3 produced with the three different fusion tags.

**Fig. S2 Mass spectrometry analysis of intact recombinant SnTox3$^{21\text{-}230}$ protein prior to and following crystallisation.** The deconvoluted spectra of protein prior to crystallisation (top panel) and within the crystallisation drop (bottom panel) show the presence of an ~24 kDa species (SnTox3$^{21\text{-}230}$). The protein within the crystallisation drop shows the presence of a ~18 kDa species, which is not present in the sample prior to crystallisation. Peaks of interest are indicated by red arrows.

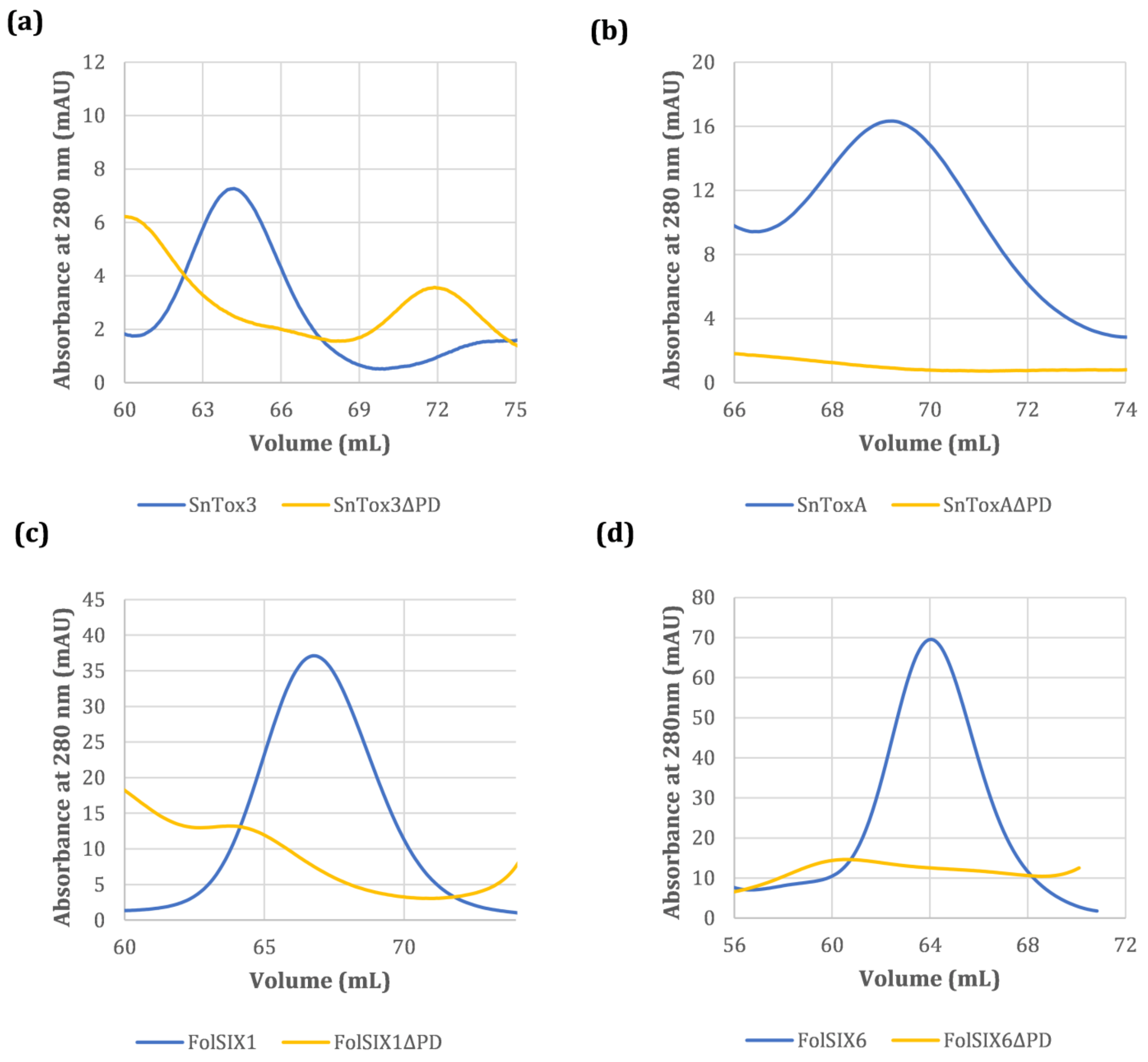


**Fig. S3 Pro-domain removal results in a significant reduction in protein solubility.** Size exclusion chromatography profiles for **(a)** SnTox3$^{21\text{-}230}$ and SnTox3$^{73\text{-}230}$(ΔPD); **(b)** SnToxA$^{17\text{-}178}$ and SnTox3$^{61\text{-}178}$(ΔPD); **(c)** FolSIX1$^{22\text{-}284}$ and FolSIX1$^{96\text{-}28}$(ΔPD); **(d)** FolSIX6$^{17\text{-}225}$ and FolSIX6$^{62\text{-}225}$(ΔPD). All proteins were produced with as an N-terminal 6xHis-GB1 fusion.

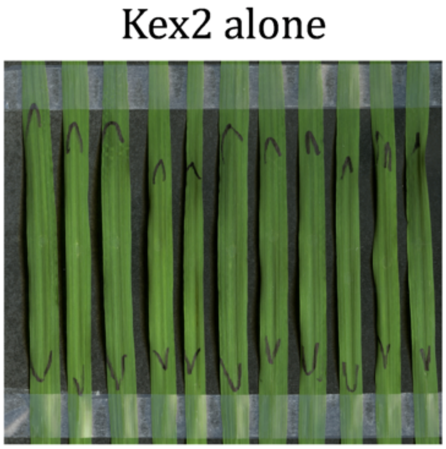


**Fig. S4 Infiltration of Kex2 alone into Corack wheat does not produce necrosis.** Recombinant Kex2 protease was infiltrated at the same concentration used to cleave SnTox3$^{21-230}$. Leaves were harvested and imaged at 3 days post-infiltration.

(a)

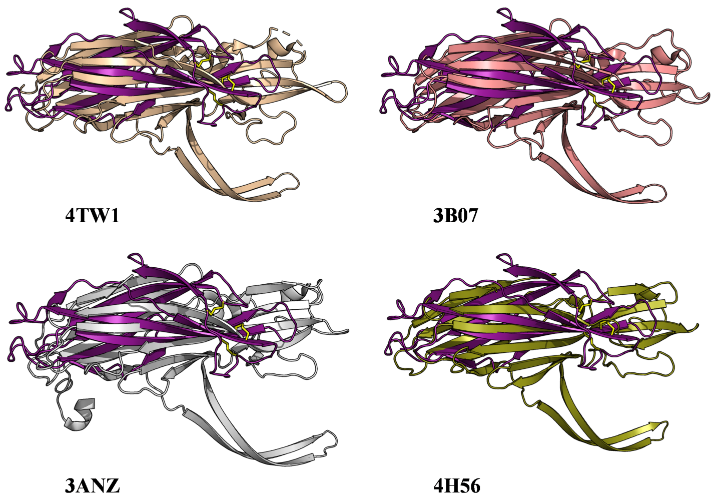


(b)

| PDB code | Z-score | RMSD (Å) | Length aligned | Number of residues | Sequence identity (%) | PDB Description |
|---|---|---|---|---|---|---|
| 4TW1 | 7.1 | 3.7 | 107 | 287 | 5 | Bi-component toxin LukGH from *Staphylococcus aureus* |
| 3B07 | 6.1 | 3.5 | 109 | 284 | 5 | Gamma-hemolysin from *Staphylococcus aureus* |
| 3ANZ | 5.8 | 3.6 | 101 | 295 | 9 | Alpha-hemolysin from *Staphylococcus aureus* |
| 4H56 | 5.1 | 3.2 | 88 | 267 | 8 | NetB toxin from *Clostridium perfringens* |

(c)

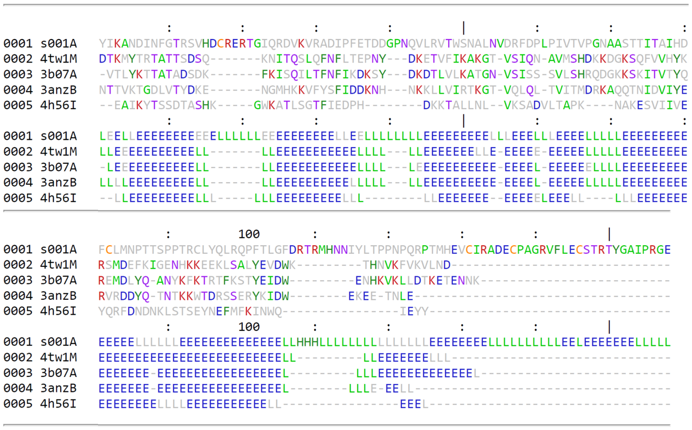


(d)

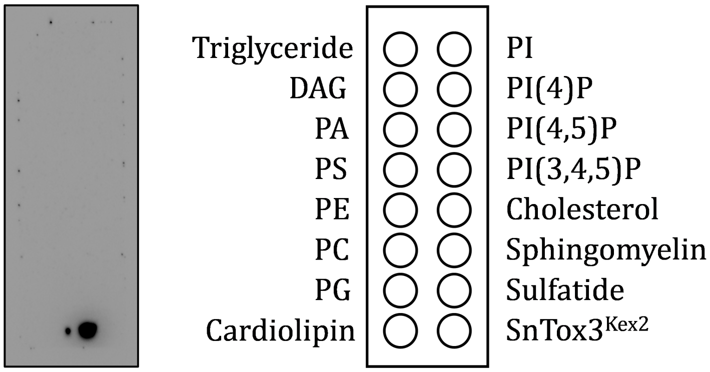

**Fig. S5 Structural-similarity search for SnTox3. (a)** Structure superimposition of SnTox3 and four unique hits showing similarity in the overall fold. The SnTox3 structure is shown in purple, and the PDB codes for the superimposed structures are indicated below each figure. **(b)** Table showing top four unique structure hits from the Dali server; they are all bacterial pore-forming toxins. **(c)** Structure-based sequence alignment showing SnTox3 has low sequence similarity with structure hits based on Dali analysis. Secondary structure alignment is shown in the lower panel of the alignment, indicating modest similarities. β-strands are denoted as E and loops as L. **(d)** Recombinant SnTox3$^{Kex2}$ was used to probe Echelon membrane lipid strips to determine if the protein could bind to membrane lipids. SnTox3$^{Kex2}$ was spotted onto the membrane (bottom right) as a control for antibody detection.

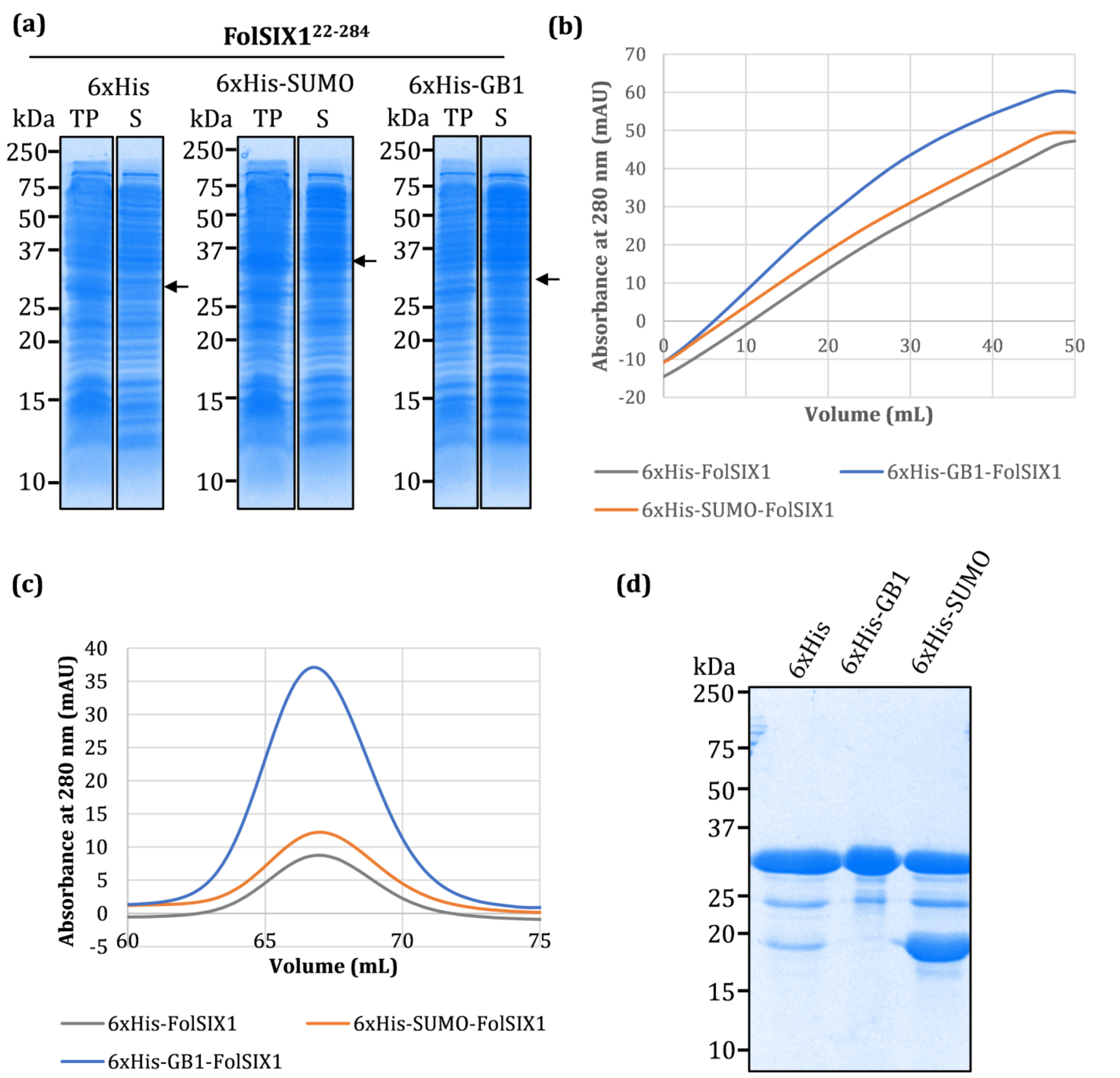


**Fig. S6 Comparison of fusion partners for FolSIX1$^{22\text{-}284}$ protein production. (a)** Coomassie-stained SDS-PAGE gel showing total protein (TP) and the soluble fraction (S) for each fusion tag (6xHis-FolSIX1, 6xHis-SUMO- FolSIX1 and 6xHis-GB1- FolSIX1). **(b)** Nickel affinity chromatography profiles for the three FolSIX1-fusion protein constructs. **(c)** Size exclusion chromatography profiles for the three FolSIX1 fusion protein constructs, following tag cleavage and removal of uncleaved protein and contaminants by a second nickel affinity chromatography step. **(d)** Coomassie-stained SDS-PAGE gel showing purified FolSIX1 produced with the three different fusion tags.

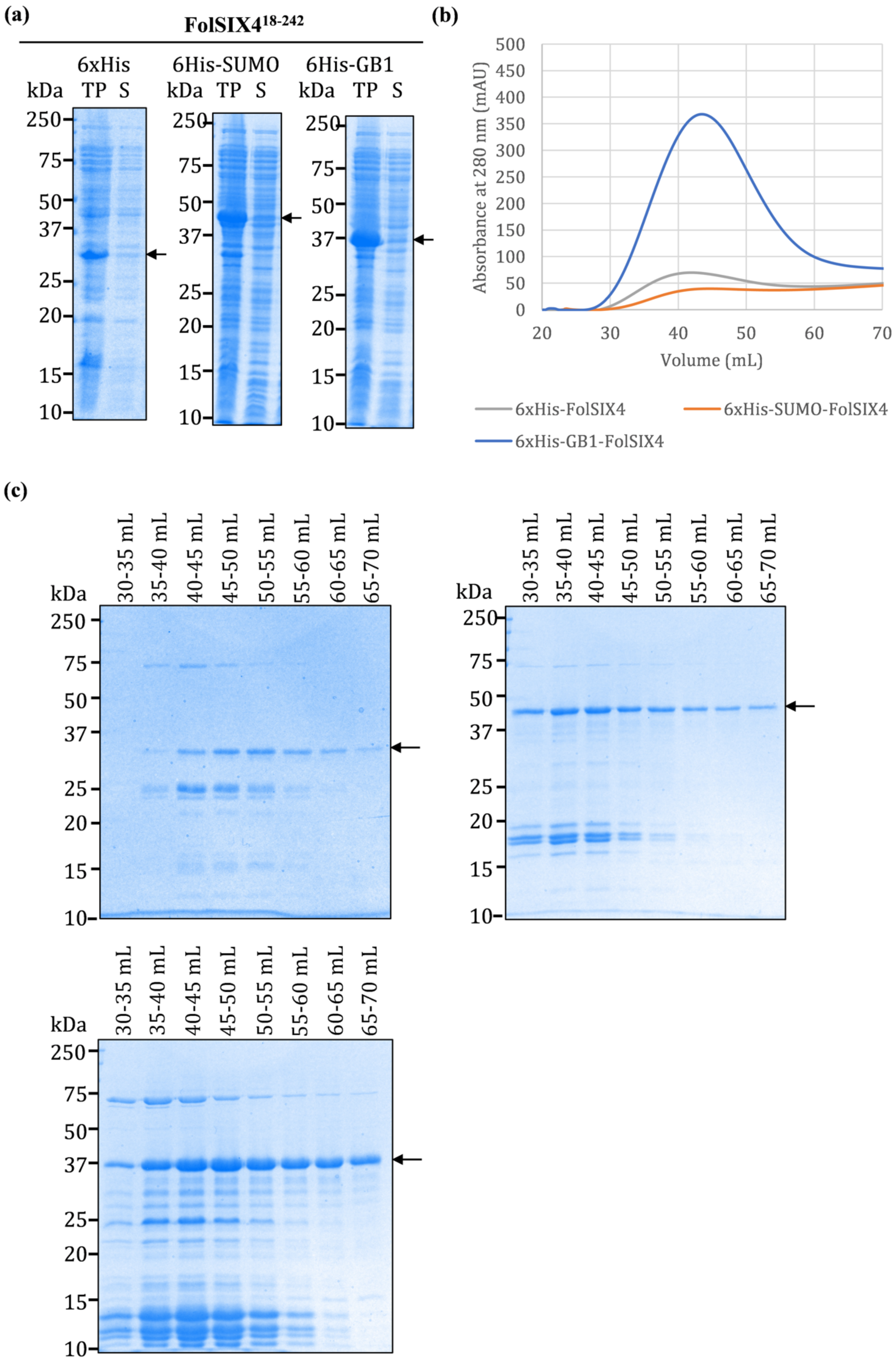
(a)
FolSIX4 18-242
6xHis
kDa TP S
6His-SUMO
kDa TP S
6His-GB1
kDa TP S
250
75
50
37
25
20
15
10
(b)
Absorbance at 280 nm (mAU)
500
450
400
350
300
250
200
150
100
50
0
20
30
40
50
60
70
Volume (mL)
6xHis-FolSIX4
6xHis-SUMO-FolSIX4
6xHis-GB1-FolSIX4
(c)
kDa
30-35 mL
35-40 mL
40-45 mL
45-50 mL
50-55 mL
55-60 mL
60-65 mL
65-70 mL
250
75
50
37
25
20
15
10

**Fig. S7 Comparison of fusion partners for FolSIX4$^{18\text{-}242}$ protein production. (a)** Coomassie-stained SDS-PAGE gel showing total protein (TP) and the soluble fraction (S) for each fusion tag (6xHis-FolSIX4, 6xHis-SUMO-FolSIX4 and 6xHis-GB1-FolSIX4). **(b)** Nickel affinity chromatography profiles for the three FolSIX4 fusion protein constructs **(c)** Coomassie-stained SDS-PAGE gel showing protein of interest (indicated by arrow) in equal-volume nickel affinity chromatography fractions for 6xHis-FolSIX4 (top left), 6xHis-SUMO-FolSIX4 (top right) and 6xHis-GB1-FolSIX4 (bottom left).

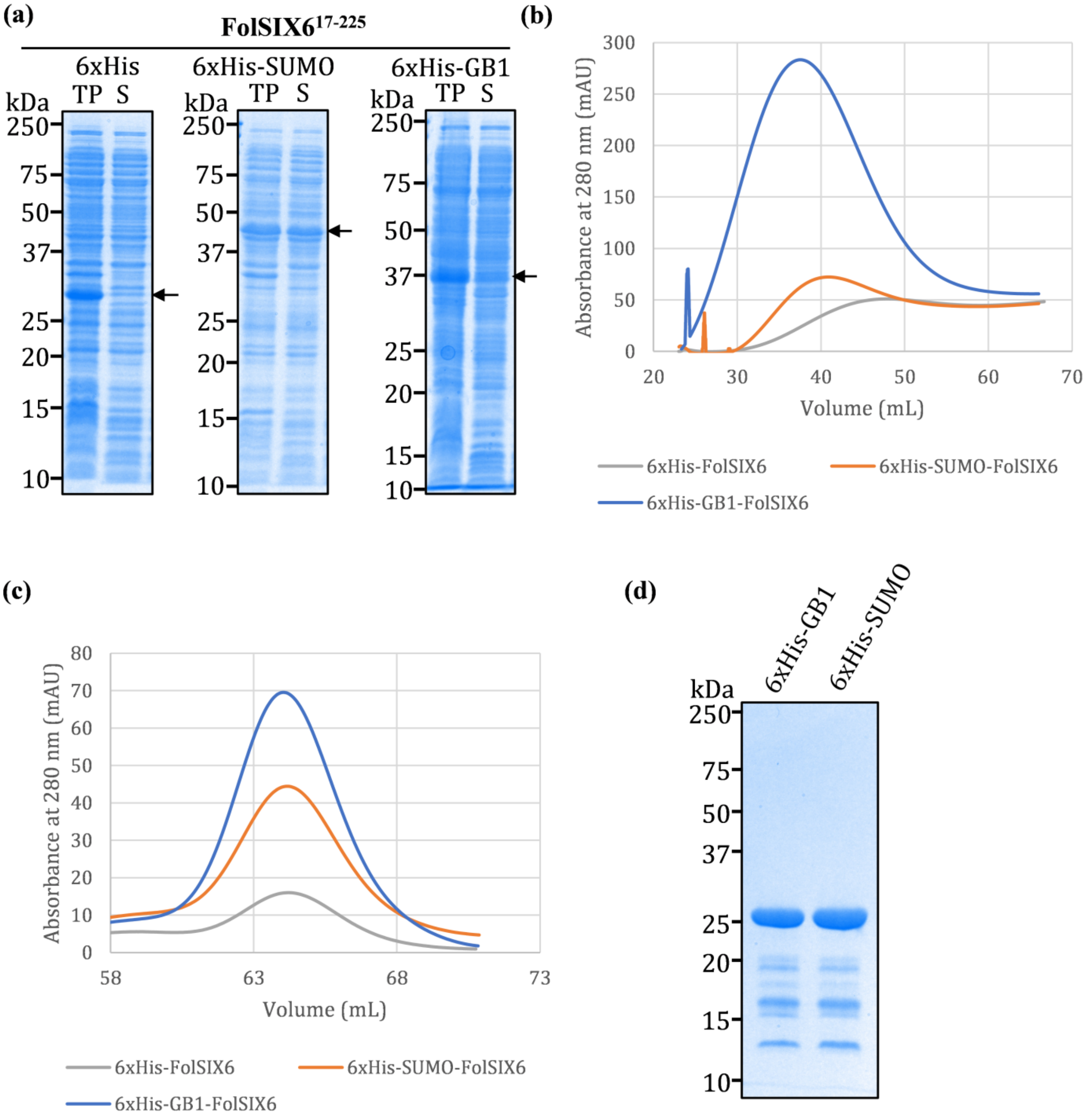
(a)
FolSIX6$^{17-225}$
6xHis
6xHis-SUMO
6xHis-GB1
TP S
kDa
250
75
50
37
25
20
15
10
(b)
Absorbance at 280 nm (mAU)
300
250
200
150
100
50
0
20 30 40 50 60 70
Volume (mL)
6xHis-FolSIX6
6xHis-SUMO-FolSIX6
6xHis-GB1-FolSIX6
(c)
Absorbance at 280 nm (mAU)
80
70
60
50
40
30
20
10
0
58 63 68 73
Volume (mL)
6xHis-FolSIX6
6xHis-SUMO-FolSIX6
6xHis-GB1-FolSIX6
(d)
6xHis-GB1
6xHis-SUMO
kDa
250
75
50
37
25
20
15
10

**Fig. S8 Comparison of fusion partners for FolSIX6$^{17\text{-}225}$ protein production. (a)** Coomassie-stained SDS-PAGE gel showing total protein (TP) and the soluble fraction (S) for each fusion tag (6xHis-FolSIX6, 6xHis-SUMO- FolSIX6 and 6xHis-GB1-FolSIX6). **(b)** Nickel affinity chromatography profiles for the three FolSIX6-fusion protein constructs. **(c)** Size exclusion chromatography profiles for the three FolSIX6-fusion protein constructs, following tag cleavage and removal of uncleaved protein and contaminants by a second nickel affinity chromatography step. **(d)** Coomassie-stained SDS-PAGE gel showing purified FolSIX6 produced with a GB1 or SUMO fusion. Protein yields for 6xHis-SIX6 were too low following concentration and were not analysed by SDS-PAGE.

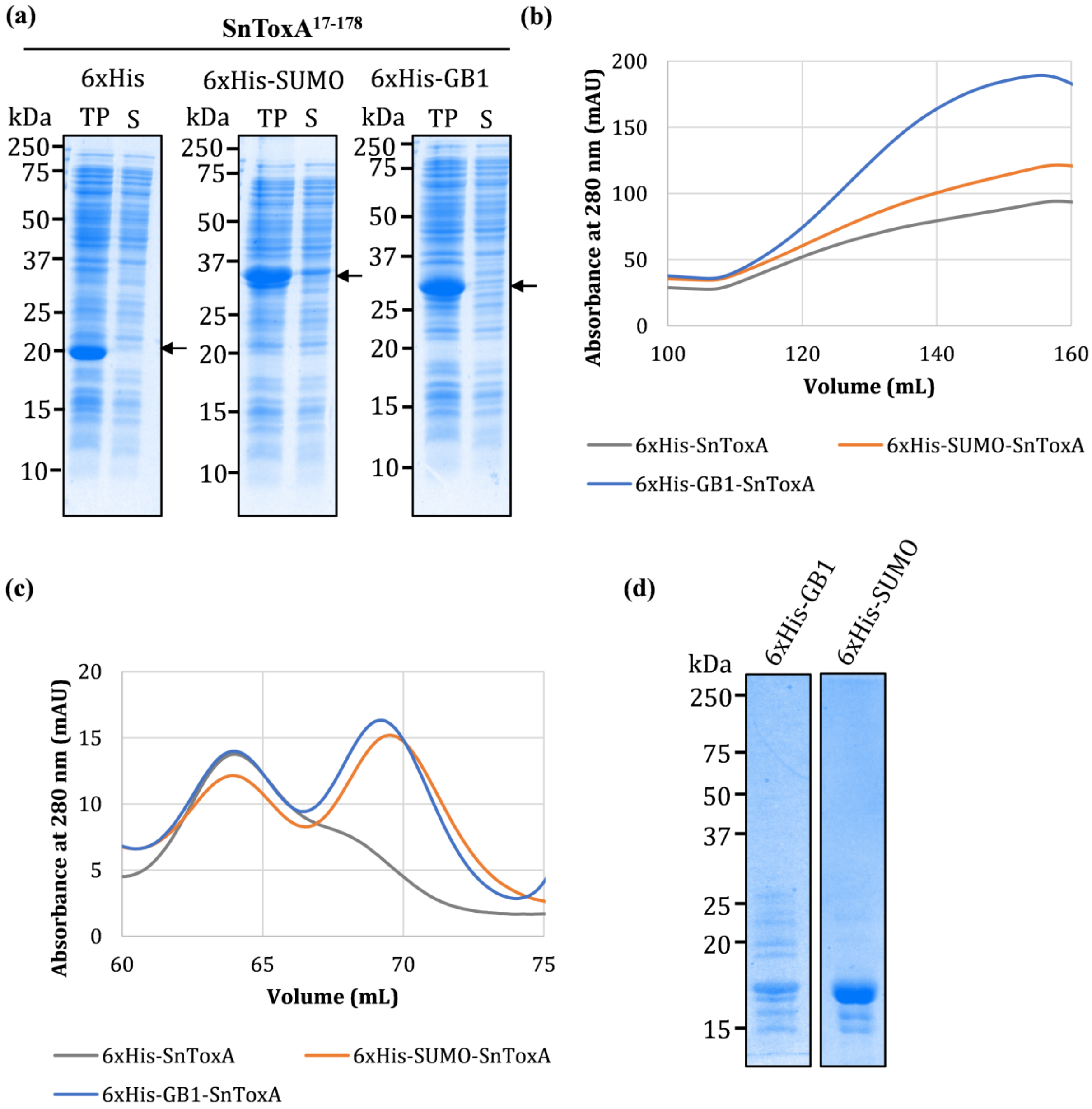
(a)
SnToxA17-178
6xHis
6xHis-SUMO
6xHis-GB1
kDa
TP
S
250
75
50
37
25
20
15
10
(b)
Absorbance at 280 nm (mAU)
200
150
100
50
0
100
120
140
160
Volume (mL)
6xHis-SnToxA
6xHis-SUMO-SnToxA
6xHis-GB1-SnToxA
(c)
Absorbance at 280 nm (mAU)
20
15
10
5
0
60
65
70
75
Volume (mL)
6xHis-SnToxA
6xHis-SUMO-SnToxA
6xHis-GB1-SnToxA
(d)
6xHis-GB1
6xHis-SUMO
kDa
250
75
50
37
25
20
15

**Fig. S9 Comparison of fusion partners for SnToxA$^{17\text{-}178}$ protein production. (a)** Coomassie-stained SDS-PAGE gel showing total protein (TP) and the soluble fraction (S) for each fusion tag (6xHis-SnToxA, 6xHis-SUMO-SnToxA and 6xHis-GB1-SnToxA). **(b)** Nickel affinity chromatography profiles for each of the three SnToxA-fusion protein constructs. **(c)** Size exclusion chromatography profiles for the three SnToxA-fusion protein constructs, following tag cleavage and removal of uncleaved protein and contaminants by a second nickel affinity chromatography step. **(d)** Coomassie-stained SDS-PAGE gel showing purity of purified SnToxA produced with a GB1 or SUMO fusion. Protein yields for 6xHis-SnToxA were low following SEC and were not analysed by SDS-PAGE.

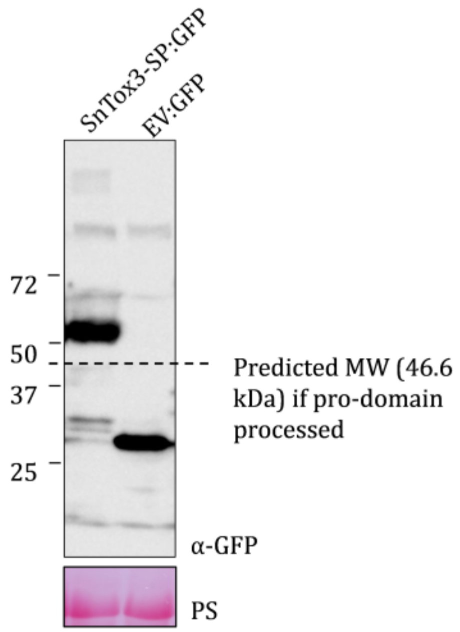


**Fig. S10 Western blot analysis of SnTox3 expression *in planta*.** SnTox3-SP:GFP (52.6 kDa) and EV:GFP (26.9 kDa) were transiently expressed in *Nicotiana benthamiana*. Leaf discs expressing the recombinant proteins were harvested at 2 days post infiltration and ground in 1.5 ml microcentrifuge tubes, followed by western blot analysis. Polyclonal rabbit anti-GFP (Santa Cruz Biotech) and goat anti-rabbit-HRP (Sigma) were used as the primary and secondary antibodies, respectively. Ponceau S (PS) staining shows protein loading.

**Table S1 Primers used in this study.**

| Primer name | Primer sequence (5'→3') |
|---|---|
| SnTox3$^{21-230}$ LIC Fw | TTATCCACTTCCAATGTTATTATTCACCACGCGGGATCGCAC |
| SnTox3$^{230}$ LIC Rv | TTATCCACTTCCAATGTTATTATTCACCACGCGGGATCGCAC |
| SnTox3$^{21-230}$ GG Fw | TAGGTCTCCAATGCTGGAACCGCGCGGTCCTGG |
| SnTox3$^{73-230}$ GG Fw | TAGGTCTCCAATGTACATCAAAGCCAACGACATCAA |
| SnTox3$^{230}$ GG Rv | ACGGTCTCCAAGATTCCCCCCGCGGGATAGCTC |
| SnToxA$^{17-178}$ GG Fw | TAGGTCTCCAATGGCCCCAACGCCTGAAGC |
| SnToxA$^{61-178}$ GG Fw | TAGGTCTCCAATGCAGGGAAGCTGCATGTCAATCA |
| SnToxA$^{178}$ GG Rv | ACGGTCTCCAAGAATTTTCTAGCTGCATTCTCCAATTTTC |
| Fol SIX1$^{22-284}$ GG Fw | TAGGTCTCCAATGCAAGAGGCTGCGGTTCGGGA |
| Fol SIX1$^{96-284}$ GG Fw | TAGGTCTCCAATGGAGCCTTTCGGGGAGGAGTCTC |
| Fol SIX1$^{284}$ GG Rv | ACGGTCTCCAAGAGTGTGGGCTGGTATATCCAAACGCC |
| Fol SIX4$^{18-242}$ GG Fw | TAGGTCTCCAATGCTTCCAAAGGGGGAGGAGGGTG |
| Fol SIX4$^{59-242}$ GG Fw | TAGGTCTCCAATGTCTGCTCATACCGAGTCTGTTTGTGT |
| Fol SIX4$^{242}$ GG Rv | ACGGTCTCCAAGAAGCTAAGTTAAGTGTACCTTGAATGCGA |
| Fol SIX6$^{17-225}$ GG Fw | TAGGTCTCCAATGGGTCCCTTAGCCCAAACAGAATCCG |
| Fol SIX6$^{62-225}$ GG Fw | TAGGTCTCCAATGGTCAGTACCTGTCCTGCGGGTC |
| Fol SIX6$^{225}$ GG Rv | ACGGTCTCCAAGATTCCCAGCTCCACGTATAGGCGTTAAG |

**Table S2** Constructs used for expression studies. All constructs were cloned using Golden Gate cloning into the pOPIN_F3_RFP_AmpR backbone, except for 6xHis-SUMO-TEV-SnTox321-230, which was cloned into the pET His6 Sumo TEV LIC cloning vector (2S-T) using ligation independent cloning.

| Construct name | N-terminal tag | Insert or PCR product | Primers |
|---|---|---|---|
| 6xHis-SUMO-TEV-SnTox3$^{21-230}$ | 6xHis-SUMO-TEV | SnTox3$^{21-230}$ | SnTox3$^{21-230}$ LIC Fw & SnTox3$^{230}$ LIC Rv |
| 6xHis-3C-SnTox3$^{21-230}$ | 6xHis-3C | SnTox3$^{21-230}$ | SnTox3$^{21-230}$ GG Fw & SnTox3$^{230}$ GG Rv |
| 6xHis-SUMO-3C-SnTox3$^{21-230}$ | 6xHis-SUMO-3C | SnTox3$^{21-230}$ | SnTox3$^{21-230}$ GG Fw & SnTox3$^{230}$ GG Rv |
| 6xHis-GB1-3C-SnTox3$^{21-230}$ | 6xHis-GB1-3C | SnTox3$^{21-230}$ | SnTox3$^{21-230}$ GG Fw & SnTox3$^{230}$ GG Rv |
| 6xHis-GB1-3C-SnTox3$^{73-230}$ | 6xHis-GB1-3C | SnTox3$^{73-230}$ | SnTox3$^{72-230}$ GG Fw & SnTox3$^{230}$ GG Rv |
| 6xHis-3C-SnToxA$^{17-178}$ | 6xHis-3C | SnToxA$^{17-178}$ | SnToxA$^{17-178}$ GG Fw & SnToxA$^{178}$ GG Rv |
| 6xHis-SUMO-3C-SnToxA$^{17-178}$ | 6xHis-SUMO-3C | SnToxA$^{17-178}$ | SnToxA$^{17-178}$ GG Fw & SnToxA$^{178}$ GG Rv |
| 6xHis-GB1-3C-SnToxA$^{17-178}$ | 6xHis-GB1-3C | SnToxA$^{17-178}$ | SnToxA$^{17-178}$ GG Fw & SnToxA$^{178}$ GG Rv |
| 6xHis-GB1-3C-SnToxA$^{61-178}$ | 6xHis-GB1-3C | SnToxA$^{61-178}$ | SnToxA$^{61-178}$ GG Fw & SnToxA$^{178}$ GG Rv |
| 6xHis-3C-FolSIX1$^{22-284}$ | 6xHis-3C | FolSIX1$^{22-284}$ | FolSIX1$^{22-284}$ GG Fw & FolSIX1$^{284}$ GG Rv |
| 6xHis-SUMO-3C-FolSIX1$^{22-284}$ | 6xHis-SUMO-3C | FolSIX1$^{22-284}$ | FolSIX1$^{22-284}$ GG Fw & FolSIX1$^{284}$ GG Rv |
| 6xHis-GB1-3C-FolSIX1$^{22-284}$ | 6xHis-GB1-3C | FolSIX1$^{22-284}$ | FolSIX1$^{22-284}$ GG Fw & FolSIX1$^{284}$ GG Rv |
| 6xHis-GB1-3C-FolSIX1$^{96-284}$ | 6xHis-GB1-3C | FolSIX1$^{96-284}$ | FolSIX1$^{62-284}$ GG Fw & FolSIX1$^{284}$ GG Rv |
| 6xHis-3C-FolSIX6$^{17-225}$ | 6xHis-3C | FolSIX6$^{17-225}$ | FolSIX6$^{17-225}$ GG Fw & FolSIX6$^{225}$ GG Rv |
| 6xHis-SUMO-3C-FolSIX6$^{17-225}$ | 6xHis-SUMO-3C | FolSIX6$^{17-225}$ | FolSIX6$^{17-225}$ GG Fw & FolSIX6$^{225}$ GG Rv |
| 6xHis-GB1-3C-FolSIX6$^{17-225}$ | 6xHis-GB1-3C | FolSIX6$^{17-225}$ | FolSIX6$^{17-225}$ GG Fw & FolSIX6$^{225}$ GG Rv |
| 6xHis-GB1-3C-FolSIX6$^{62-225}$ | 6xHis-GB1-3C | FolSIX6$^{62-225}$ | FolSIX6$^{62-225}$ GG Fw & FolSIX6$^{225}$ GG Rv |
| 6xHis-3C-FolSIX4$^{18-242}$ | 6xHis-3C | FolSIX4$^{18-242}$ | FolSIX4$^{18-242}$ GG Fw & FolSIX4$^{242}$ GG Rv |
| 6xHis-SUMO-3C-FolSIX4$^{18-242}$ | 6xHis-SUMO-3C | FolSIX4$^{18-242}$ | FolSIX4$^{18-242}$ GG Fw & FolSIX4$^{242}$ GG Rv |

| | | | |
|---|---|---|---|
| 6xHis-GB1-3C-FolSIX4$^{18-242}$ | 6xHis-GB1-3C | FolSIX4$^{18-242}$ | FolSIX4$^{18-242}$ GG Fw & FolSIX4$^{242}$ GG Rv |
| 6xHis-GB1-3C-FolSIX4$^{59-242}$ | 6xHis-GB1-3C | FolSIX4$^{59-242}$ | FolSIX4$^{59-242}$ GG Fw & FolSIX4$^{242}$ GG Rv |

### Table S3 X-ray data collection, structure solution and refinement statistics for SnTox3.

| | SnTox3 Bromide soak (SAD) | SnTox3 Native (MR) |
|---|---|---|
| **Data collection** | | |
| Detector | ADSC Quantum 315r CCD | ADSC Quantum 315r CCD |
| Wavelength (Å) | 0.9116 | 0.9537 |
| Crystal-to-detector distance (mm) | 220 | 120 |
| Space group | C121 | C121 |
| *a*, *b*, *c* (Å) | 90.57 29.69 56.36 | 99.87 29.91 56.05 |
| a, b, g (°) | 90.0 101.72 90.0 | 90.0 102.12 90.0 |
| Average mosaicity (°)[b] | 0.16 | 0.85 |
| Resolution (Å) | 19.48-2.1 (2.10-2.04) | 28.6-1.36 (1.37-1.35) |
| Total no. of reflections | 155810 (11295) | 254407 (11339) |
| No. of unique reflections | 10575 (820) | 35938 (1759) |
| Completeness (%) | 99.9 (99.7) | 98.7 (99.1) |
| Multiplicity | 14.7 (13.8) | 7.1 (6.4) |
| Anomalous completeness | 99.8 (99.8) | - |
| Anomalous multiplicity | 7.7 (7.1) | - |
| Mean *I* /s(*I*) | 18.0 (4.8) | 15.6 (3.5) |
| *R*meas (%)[c] | 14.4 (63.7) | 7.3 (47.4) |
| *R*pim (%)[d] | 5.2 (23.6) | 7.1 (47.4) |
| $CC_{1/2}$[b] | 0.998 (0.950) | 0.999 (0.914) |
| Matthews coefficient ($Å^3$ $Da^{-1}$)[e] | 2.29 | 2.3 |
| **Refinement** | | |
| Resolution range (Å) | | 26.02-1.35 |
| $R_{work}$ (%)[g] | | 17.3 |
| $R_{free}$ (%)[h] | | 19.9 |
| No. of non-H atoms | | |
| Total | | 1410 |
| Non-solvent | | 1253 |
| Water | | 157 |
| Average *B*-factor ($Å^2$) | | 17.0 |
| R.m.s.d. from ideal geometry | | |
| Bond lengths (Å) | | 0.50 |
| Bond angles (°) | | 0.58 |
| Ramachandran plot, residues in (%)[i] | | |
| Favoured regions | | 98.08 |
| Additionally allowed regions | | 1.92 |
| Outlier regions | | 0 |

[a] The values in parentheses are for the highest-resolution shell.
[b] Calculated with AIMLESS (Evans & Murshudov, 2013).
[c] $Rmeas = \Sigma_{hkl}\{N(hkl)/[N(hkl)-1]\}^{1/2} \Sigma i|Ii(hkl)-<I(hkl)>|/ \Sigma hkl \Sigma i Ii(hkl)$, where *Ii*(*hkl*) is the intensity of the *i*th measurement of an equivalent reflection with indices *hkl*.
[d] $Rpim = \Sigma_{hkl}\{1/[N(hkl)-1]\}^{1/2} \Sigma i|Ii(hkl)-<I(hkl)>|/ \Sigma hkl \Sigma i Ii(hkl)$.
[e] Calculated with MATTHEWS_COEF within the CCP4 suite (Winn *et al.*, 2011).
[f] Generated by Crank pipeline (Pannu *et al.,* 2011) in the CCP4 suite (Winn *et al.*, 2011).
[g] $R_{work} = \Sigma_{hkl} ||F_{obs}|-|F_{calc}||/\Sigma_{hkl} |F_{obs}|$, where $F_{obs}$ and $F_{calc}$ are the observed and calculated structure factor amplitudes.
[h] $R_{free}$ is equivalent to $R_{work}$ but calculated with reflections (5%) omitted from the refinement process.
[i] Calculated with MolProbity (Davis *et al.*, 2004).

**Table S4 Effectors used for protein production analysis, and obtained yields.**

| Effector | Organism | Accession number | No. of cysteines | Yield (mg/L of culture) | | |
|---|---|---|---|---|---|---|
| | | | | 6xHis | SUMO | GB1 |
| SnTox3$^{21\text{-}230}$ | *Parastagonospora nodorum* | ACR78113.1 | 6 | 0.05 | 0.54 | 0.83 |
| SnToxA$^{17\text{-}178}$ | *Parastagonospora nodorum* | XP_001806667.1 | 2 | 0.05 | 0.11 | 0.23 |
| FolSIX1 (Avr3)$^{22\text{-}284}$ | *Fusarium oxysporum* f. sp. *lycopersici* | ACY39281.1 | 8 | 0.07 | 0.10 | 0.25 |
| FolSIX4 (Avr1)$^{18\text{-}242}$ | *Fusarium oxysporum* f. sp. *lycopersici* | ACY39284.1 | 6 | 0.05 | 2.00 | - |
| FolSIX6$^{17\text{-}225}$ | *Fusarium oxysporum* f. sp. *lycopersici* | XP_018253967.1 | 8 | 0.10 | 0.15 | 0.27 |

**Table S5** All possible Kex2 cleavage motifs present in putative Kex2-processed pro-domain (K2PP) fungal effectors identified via in silico analysis. Signal peptides were predicted using SignalP 5.0 and are shown in bold. Cysteine residues are highlighted in yellow. Putative Kex2 cleavage motifs identified using our in silico analysis are shown in cyan, and alternative putative Kex2 cleavage motifs not identified are shown in green.

| **Predicted K2PP Effector** | **Species** | **Sequence** |
|---|---|---|
| AvrSr35 | *Puccinia graminis* | **MSHHFGLRKIKLLILLFLQVHGSQC**AMRNFAADRVHGVESVISGSKSSSNPMALSKSMDKPDTSDLVDSNVQAKNDGSRYEEDFTAKYSEQVDHVSKILKEIEEQEPGTIIIDHKAFPIQDKSPKQVVNFPFPKKMITESNSKDIREYLASTFPFEQQSTILDSVKSIAKVQIDDRKAFDLQLKFRQENLAELKDQIILSLGANNGNQNWQKLLDYTNKLDELSNTKISPEEFIEEIQKVLYKVKLESTSTSKLYSQFNLSIQDFALQIIHSKYKSNQISQNDLLKLITEDEMLKILAKTKVLTYKMKYFDSASKMGINKYISTEMMDLDWQFSHYKTFNDALKKNKASDSSYLGWLTHGYSIKYGLSPNNERSMFFQDGRKYAELYAFSKSPHRKIIPGEHLKDLLAKINKSKGIFLDQNALLDKRIYAFHELNTLETHFPGITSSFTDDLKSNYRKKMESVSLTCQVLQEIGNIHRFIESKVPYHSSTEYGLFSIPKIFSIPIDYKHGEKENLVSYVDFLYSTAHERILQDNSINQLCLDPLQESLNRIKSNIPVFFNLASHSSPIKPSNVHEGKL |
| FGB1 | *Piriformospora indica* | **MKFTTVFVTLAIFASSAFA**AAVADDETAVIARARSEDAAGSVDLLKRACNYNTPCKGYYWSKGLHCGDGNYGCVYGHVYQVGSDTNVCDYGVRTSCQKCGKLSC |
| FGSG_10999 | *Fusarium graminearum* | **MVSFKSLLVAVSALTGALA**RPFDFLDERDDGNATSVLEARQVTGNSEGYHNGYFYSWWSDGGGYAQYRMGEGSHYQVDWRNTGNFVGGKGWNPGTGRTINYGGSFNPQGNGYLCVYGWTRGPLVEYYVIESYGSYNPGSQAQHRGTVYTDGDTYDLYMSTRYQQPSIDGVQTFNQYWSIRRNKRTSGSVNMQNHFNAWRSAGMNLGNHYYQILATEGYQSSGSSSIYVQTS |
| SAD1 | *Sporisorium reilianum* | **MNLLPFKLVLALSLLLLTLSGCHC**ASGSQSGGGNHNVEVDEQAAQDLRARVAHLERHFKLGPNWFGETRGPHVAQEVARSQLRLTSAKFMVHHDPADYRFNLAYTPVSYRMPDSADWKHGILMLRVVPHQPTQVLDLVDFPEGLAQREVEGYWGRFYGAVDKTREDILREYGEPALHLSV |

| | | |
|---|---|---|
| MISSP7 | *Laccaria bicolor* | **MKLLVPLNLLIAALAISPALS**SPVPGEVGLVERGPIPNAVFRRVPEPNFFKDLLR<br>ALGQASQGGDLHR |
| Mig2-5 | *Ustilago maydis* | **MFFHKPLLLAASLSLFLCLFPFASA**GRIDRGQLRFSKRSPAPSSEKKDSALGML<br>HAKWSANNKRAPPSIPPTGYTTITSTLKILPEDAMPLGECYCPRTLDEESQGISGA<br>CAEKFVASAKKTAKCTFLTSTEATCLEEGGKANLSDGDTAWWQMAFCKRCQD<br>AGGISNALYPSCAAQMPQD |
| Mig2-4 | *Ustilago maydis* | **MFFHKPLFLAASLSLFLCLFTFASA**GRANGDQQYLRKRQSETSIIGRTEGPSRPV<br>PFTSSPDLVVITSKLEILGKESSALGQCSCVRTLDDQKQNVSGTCTSRFVDSAKST<br>ATCLFGTLTDATCFEKGGKLDLSDGDKVWWQKAFCKRCKAAGGKSHPFYACP |
| Mig2-1 | *Ustilago maydis* | **MFSYNSFLLATAILCHFFCACRFTSA**VQINLNKRGLHRRSIGTQLGSVLEHRPV<br>RVEARSDITPGIVAGNDGHSSGENWILITSSLKKRGDNHQFLGLCICDPSSADQKQ<br>STTGSCTSKFQASAEKTADCKFTAKTILDCFEEGGKQDMSAGDNVWWQMAFCK<br>RCKEAGGTSHPSYACP |
| Mig2-2 | *Ustilago maydis* | **MFFHKSLLLATASLSLFFCEFSFTSA**SPINVEQRGLQRQSVETSLDVRDDLILGR<br>GHQRKRDLRFSDGDESHVDKSADARSDSGRLGKLDSRHGGVDLNIIGHPRLDRR<br>DKCISDCPHAGDPNGGPDPGHPHVSKRDNGIDADSPEQISRPGWPRNGIPYPPWG<br>HTHVVQRDHGLGWQGPGGTFRPGWPPRDIPPPPMTNCQEVGQRPPASSKIGRDS<br>IEKRNDISHALRRRADGVSGDVSTSDTTIATSKPDLSKIGIGKGGAHRGVPRFHGG<br>EAPLPSFPSPPSPPHHPPISSVGAVPDSVHDQNVRFSGPRPFERLPTRHLGQFEKRS<br>PLQLSDDEGSLPDALHGAWNDSEHEVSSGSNPNSNVGGSGKNVDPLTTQLLQNP<br>KQPYLRGCTRFGECNGAHELSEPADRIGETDSLVEEPPSFAPRHKQDLKHHSLEL<br>TRRYYGENGDISAREHEPKLLEARSKMMPTLFTRDHVDQTSQKPVLITSSVQHHP<br>ADIQVFGICTCNGSTEVQKPDTKVNCLSEFLNSAKKTAKCSFTASDKVTCVEEGG<br>KDDMSGGDNVWWQIAFCKRCQDAGGTSDPKFKCH |
| Rsp3 | *Ustilago maydis* | **MKFQASFVTLLVVVSAAISVA**DAKAPMANKRPSNLNRRDEAAALQKRGSLGG<br>FANRFFRRDGGAYGDNQDADAGGDTANPDSQNGNQSGDQPPPPDTDDGSVQQ<br>GYGNDERHQGQPPPSQPQPQPQPQPQPQTQPQPQPQPQPQGQDSQQQPKTDQ<br>QKKDEEKQKQDEQKRQQEEAQRQQKEQERQKKDEEKAKQQSQGGSQNGNNQP<br>IGSGNGGQPVAPNQNQETTIPDNSQGQGSGPDNCPKFPKKGGGETDHAPCFSKD<br>ANRKDICSAMKQFGIDTKAVGCSDDECSEDAKDFSNHACNKKKQCDFNWDNSC<br>DFYLHMGFTNVAKFGCGAPNGGDDGDSSLPSDDSSAGQQPRDNNPGQQPGNNN<br>PGQQPGNNNPGQQPGDNNPGQQAGKDANEDEECDDEGDDKGDNGAAGGDKN<br>PNGDNGAAGGDKNPNGDNSSEGDEPVPYGDETGDADNGAAGGDKNPSGDKNP |

| | | |
|---|---|---|
| | | NGDKNPSGDKNPNGDNGAAGGDKNPNGDNGAAGGDKNPNGDNGAAGGDKNP<br>NGDNSSEGDEPVPYGDETGDADNGAAGGDKNPNGDNSSEGDEPVPYGDETGDA<br>DNGAAGGDKNPSGDKNPNGDKNPNGDNGAAGGDKNPNGDNGAAGGDKNPNG<br>DNSSEGDEPVPYGDETGDADNGAAGGDKNPSGDKNPNGDKNPSGDKNPNGDN<br>GAAGGDKNPSGDKNPNGDNGAAGGDKNPNGDNGAAGGDKNPNGDNGAAGG<br>DKNPNGDNGAAGGDKNPNGDNSSEGDEPVPYGDETGDADNGAAGGDKNPSGD<br>NGAAGGDKNPNGDNGAAGGDKNPNGDNSSEGDEPVPYGDETGDADNGAAGD<br>KGANKPDDCKKNKDASAGGDTQDSKDAQGDSKAPQP |
| eff1-1 | *Ustilago maydis* | **MGLRLRILALLLAVSVVAA**WPYPSGNHPDNADGARRGEQSPLPGFSVAHPSSP<br>QNEFDPDLLRSLLAKIESEASSHPHVAFQGDGHYYPNVNQPSSLYQASQHGFGSL<br>SRDKGQVGSSQARESFSDTQGLSSSAEPAFKYRKTVQRVADALRQFQANNRLEG<br>AQALRDIIQRHPQLDTTSGSQSQGFRNNFGAAPSVPSRESQGGTIHHGPGDTKVV<br>DIPSDRIEAQTTAEESHESQPLAGHGDGRYRSLPASRADELRKIPTTMLALPPPSR<br>TRYIYDQVDDPAIRDNINNQVFAGKLVWIDRARIPASRIVSSRKQMFRPHRVLPM<br>TNFPEIHLSDGKGSIKDVRFTFHGGGSYRLMSWPEGYNLVEGQNYLAFWGIPEG<br>RQTGRPMLMQNYGYAFLPPQHRLEVNEHLWALKKEIADKASEATWMHA |
| Tin2 | *Ustilago maydis* | **MNRLQSYTRVFIALVLFSMVCNCFA**TGGFDYENLASSSSSGHHDYRLESSPM<br>GRFQTKDYANLLNSRLKDMGWGSEPLAKGLLVPREDLESFVSSVQDVLKTESG<br>QKGLLYLGNTPDSRKVHAVLLKGAQSRSPNIAIISTPKVWRSLNQKIQLHTFAQID<br>NSNLFDLHQLLQHDFNYSQLDSLARNALPKTPQSEFDNLLPRFPL |
| Pit2 | *Ustilago maydis* | **MLFRSAFVLLIVAFASACLVQHVQA**IPVRRSLSTDASMSSAAGKLNRRWWFGF<br>TGSLGKEPDNGQVQIKIIPDALIIKNPPANKDDLNKLIENLKRKHPRFKTVVMPTD<br>PNGDVVIWE |
| Cmu1 | *Ustilago maydis* | **MKLSVSIFVLLAVSAFGGGSA**AAVSGKSEAAEIEAGDRLDALRDQLQRYETPII<br>QTILARSALGGRAPSEQDEVRAALSRNAFEPSEVISEWLQTESGARFRSTRPLPPA<br>VEFITPVVLSRDTVLDKPVVGKGIFPIGRRPQDPTNMDEFLDTSLLSLNQSSTVDL<br>ASAVSLDVSLLHLVSARVLLGYPIALAKFDWLHDNFCHILTNTTLSKSQKLANIIQ<br>QLTDHKQEVNVLSRVEQKSKSLSHLFRNDIPYPPHTQDRILRLFQAYLIPITTQIEA<br>AAILDHANKCT |
| VdSCP7 | *Verticillium dahliae* | **MKTCVIATLVGVAMS**APAMRTSMDAPMMEMANSRPMDMDMGSSTPAMRNT<br>MNMKDASSSMGMAKRERVEMGNMAPTMKKTKGMKMGSMAPAMKTGMATR<br>STMDMDMANRNSATKGMEKNKDVDAIVAAMLMEMAHRRPVAGMVRRQSEE |

| | | |
|---|---|---|
| | | AFQGVVEECKTKLASGEVTSLDNCVLDTLGINRRQVTGDQDQLAQITQECTEKTPNGKSTRIQGVHY |
| AVR-Pik | *Magnaporthe oryzae* | **MRVTTFNTFLLTLGTVAVVNA**ETGNKYIEKRAIDLSRERDPNFFDHPGIPVPECFWFMFKNNVRQDAGTCYSSWKMDMKVGPNWVHIKSDDNCNLSGDFPPGWIVLGKKRPGF |
| AvrPi9 | *Magnaporthe oryzae* | **MQFSQILTVLFLGVSVSA**LPAGGLPGSPGSAVQRCHCPPRGSHAHGSLAAREEAPEAEGDAKISARYTCPNCHKTGKGCDDGWCQVEKTHW |
| BAS162 | *Magnaporthe oryzae* | **MRSQALLAVLYASGAIAGGGT**YIPKDPALAHRAPEKFPWSGLLNNPFMTKPVIWYQIGQIKIWTDTRDVTYVRDGTKFDTYDQCLSHCLKYSVFNKSKARSSGSQGQTSEYRGRRMHH |
| MoCDIP1 | *Magnaporthe oryzae* | **MARFTSLIAAVLATIGAVKA**QCGAGNPDARVTGSGNNFQAVRGSNTVYSGSDYRAAIQAALDSIGSGQRVAVIASGSIGANTISISSGKTFEGCGTINVGNRNGRGAIESLNTQGVKIPYLTMTGNPYFGLRFYGTRDLTLGQITMNLSGGLGIRFDRDNPDNFNVRMGTITVTGAGSHAVETWNIDGLVIDRVIARNVGESGLLVQKTRNAQIGIVDGNNVGTGTGYATLRFANNNGQNPNGNYNTNIWVDQVISRGGGRGVFCVSQSGAAVIRTVDLAKNGNNAILIENCYNLSIRGGTVNGGGEVRVAARSEFPNNRDLWITLRVDNTSVRESPCGTNVNWSLTGNGQRALC |
| MoHEG13 | *Magnaporthe oryzae* | **MLSLKTLLLVSVSCLAQLAAA**HPLDNGGLEPRQQKICSPNGHTDGDLTNPNSLCTLCNSGCQRVAGQPACCS |
| Lug6 | *Magnaporthe oryzae* | **MQFSTIQLFALMAVGAMA**NTVAPADVAAVDAPRAPLLVRRNCEGKNTQAECERFKWLTGCTWLRGSKYCSST |
| AVR-Pii | *Magnaporthe oryzae* | **MQLSKITFAIALYAIGIAA**LPTPASLNGNTEVATISDVKLEARSDTTYHKCSKCGYGSDDSDAYFNHKCN |
| BAS3 | *Magnaporthe oryzae* | **MQFSTVSFAIFAILPAMVAA**MPAETSPVPKPALPVFEELCPDAERQKCAESTDNLKRCLQINGASICVIDCGSQTTCRTQCKQQLKNEKANGFCTVGDNPCICNLNGAANSAH |
| AvrLm1 | *Leptosphaeria maculans* | **MVQFKTIFLSTALAALFSTGSS**SPATKNNVNQPLDNISRRSEWKSVQISPVKEHSAKTADNTENNHNLEKRVFTSPHMKRTFTLALENTFYAMAWLIDFSFSDDGEPHFSYKLQSFNHEDNPPKILADVVVPLITTSYNSANQYRGKANVELLCHLAKEYVHVYFSVEVFASGASFVIGKIIDYPTVYVNNQFRKVVKFDIAGAI |
| Zt6 | *Zymoseptoria tritici* | **MRFSLLSSALLFATAAFS**APVAEPAELEIRQQATYCGNQYYSASQVSAAVNKGYNYYANGQQVGSGNYPHQYNNREGFSFAVSGPYQEFPILASGSTYSGGSPGPDRVVFNTRGQWGGTITHTGASNNNFVGCSGTS |

| | | |
|---|---|---|
| SIX8 | *Fusarium oxysporum f. sp. lycopersici* | **MQPLRILLLFPLAVSVAA**TPIDKSLDQAATIEETVHQPHSHDERALVERDTSGIL<br>LACITGAGSAFQAYAGCYLTAFRNDPRTLTLRMDKTRGERISNVLVILSGGALSH<br>AVEEVVQIAPGAVRNLATLGASTVQFLHNFR |
| SIX2 | *Fusarium oxysporum f. sp. lycopersici* | **MLFKIAWVSLFTTWAISVAA**NPAGDSLPDDAHLPDRRLSPSEVQALKKAQIYPP<br>GYIHKRVTFGEGKDAVEVPIVEADVEMLLNNEKGVKARSLAPRGSCFSFPTPAR<br>GSCMIDYCWRDDNGVIYSRGITITGSNGASNPTSMRSNDPANLSLNSVFNDGYN<br>GWFPHGHACSNSDTQIYTNHRLLQGVNGVAYVDHVRCENCNFRNVNCLSDVLK<br>NNLIAYSNGVASQSRCT |
| SIX3 | *Fusarium oxysporum f. sp. lycopersici* | **MRFLLLIAMSMTWVCSIAG**LPVEDADSSVGQLQGRGNPYCVFPGRRTSSTSFT<br>TSFSTEPLGYARMLHRDPPYERAGNSGLNHRIYERSRVGGLRTVIDVAPPDGHQA<br>IANYEIEVRRIPVATPNAAGDCFHTARLSTGSRGPATISWDADASYTYYLTISED |
| SIX6 | *Fusarium oxysporum f. sp. lycopersici* | **MKLALIASILAAGCVA**GPLAQTESESADVAEHTINYIDIAPEEFEPPKANLSSLVS<br>RDTLPVSTCPAGQKYDRSVCYKADKIRSFCVANPRSNREKITDTPCQPREICVQR<br>NLSNGKSFAKCIPIVDLVEWKTSANGNKEGCTTTSVNPAGYHHLGTIVYDINKNP<br>IEVDKISYFGEPGNVNEGIGGSTSYFSSDNFQFSKSRYMKTCIFSGGYGNLNAYT<br>WSWE |
| SIX4 | *Fusarium oxysporum f. sp. lycopersici* | **MNLKALVVIASVAVTSA**LPKGEEGDIIGTFNFSSSDSQPLKIHWVDTPDSSGSNL<br>VKRSAHTESVCVHAGTATGADLHWLNAICTGKSTYTVNCAPAGNKNAGSTHTG<br>TCPAGQDCFQLEQVGNFWGDREPDATCSPSNTVFDAVDDKEATHVNGKVVTRA<br>GKPGIGRKLIRLKAQVYRRDGHYGQTSRMGFFRNGKEVYHIDNVASMEPTWNF<br>DPSSDQSFSFFFTPGPNAFRIQGTLNLA |
| SIX1 | *Fusarium oxysporum f. sp. lycopersici* | **MAPYSMVLLGALSILGFGAYA**QEAAVREPQIFFNLTYTEYLDKVAASHGSPPD<br>KSDLPWNDTMGSFPGNETDDGVQTETGSSLSRRGHIVNLRKREPFGEESRNDRV<br>TQDMLQALHDLCVERFGTGYRAVSGLCYTDRRATRKIECNKPSVRERDRSVTRA<br>CPKGQECTTFNAYNFRNRHHQVTFPVCGPRIEVKDRHDIGIHTEWQGTWYPESP<br>KSPGTYDYFAQMAGTLNGYFGYDGVYSDGYKTSSHGYGHSWSCINCPRGKVTI<br>TNTYRATWAFGYTSPH |
| SnToxA | *Parastagonospora nodorum* | **MRSILVLLFSAAAVLA**APTPEADPGYEIVKLFEAANSSELDARGLSLDWTLKPR<br>GLLQERQGSCMSITINPSRPSVNNIGQVDIDSVILGRPGAIGSWELNNFVTIGLNRV<br>NANTVRVNINNTGRTNRLIITQWDNTLTRGDVYELFGDYALIQGRGSFCLNIRSD<br>SGRENWRMQLEN |
| SnTox3 | *Parastagonospora* | **MHFTKFLLVVQAAATAIALA**LEPRGPGDIQLTREEHEAIFNGSPSDWTEDPNFK<br>PDVPEQQRPATANDLSKRYIKANDINFGTRSVHDCRERTGIQRDVKVRADIPFET |

| | | |
|---|---|---|
| | *nodorum* | DDGPNQVLRVTWSNALNVDRFDPLPIVTVPGNAASTTITAIHDFCLMNPTTSPPTRCLYQLRQPFTLGFDRTRMHNNIYLTPPNPQRPTMHEVCIRADECPAGRVFLECSTRTYGAIPRGE |
